# Evidence for extensive anaerobic dechlorination and transformation of the pesticide chlordecone (C_10_Cl_10_O) by indigenous microbes in microcosms from Guadeloupe soil

**DOI:** 10.1101/650200

**Authors:** Line Lomheim, Laurent Laquitaine, Suly Rambinaising, Robert Flick, Andrei Starostine, Corine Jean-Marius, Elizabeth A. Edwards, Sarra Gaspard

## Abstract

Chlordecone (C_10_Cl_10_O) is a bishomocubane molecule, that has been used as pesticide, in many countries in Europe, America, and Africa, from the 1960’s to 1990’s. In the French West Indies, the historic use of chlordecone to control banana weevil infestations has resulted in pollution of large land areas. Although currently banned, chlordecone persists because it adsorbs strongly to soil and its complex structure is stable, particularly under aerobic conditions. A leaching model established that CLD pollution will last in French west indies soils several decades to half a millennium depending on soil type. However, abiotic chemical transformation catalyzed by reduced vitamin B12 has been shown to break down chlordecone by opening the cage structure to produce C9 polychloroindenes, and more recently these C9 polychloroindenes were also observed as products of anaerobic microbiological transformation by *Citrobacter*. To assess the potential for bioremediation, the anaerobic biotransformation of chlordecone by microbes native to soils from the French West Indies was investigated. Anaerobic microcosms were constructed from chlordecone impacted Guadeloupe soil and sludge to mimic natural attenuation and eletron donor-stimulated reductive dechlorination. Original microcosms and transfers were incubated over a period of 8 years, during which they were repeatedly amended with chlordecone and electron donor (ethanol and acetone). Using LC/MS, chlordecone and degradation products were detected in all the biologically active microcosms. Observed products in active incubations included monohydro-, dihydro- and trihydrochlordecone derivatives (C_10_Cl_10−n_O_2_H_n_, n= 1,2,3), as well as “open cage” C9 polychloroindene compounds (C_9_Cl_5−n_H_3+n_, n=0,1,2) and C10 carboxylated polychloroindene derivatives (C_10_Cl_4−n_O_2_H_4+n_, n=0−3). Products with as many as 9 chlorine atoms removed were detected. These products were not observed in sterile incubations. Chlordecone concentrations decreased in active microcosms as concentrations of products increased, indicating that anaerobic dechlorination processes have occurred. An crude estimation of partitioning coefficients between soil and water showed that carboxylated intermediates sorb poorly, and as a consequence may be flushed away while polychlorinated indenes sorb strongly to soil. Microbial community analysis in microcosms showed enrichment of anaerobic fermenting and acetogenic microbes possibly involved in anaerobic chlordecone biotransformation. It thus should be possible to stimuilate anaerobic dechlorination through donor amendment to contaminated soils, particularly as some metabolites (in particular pentachloroindene) were already detected in field samples as a result of intrinsic processes. Extensive dechlorination in the microcosms, with evidence for up to 9 Cl atoms removed from the parent molecule is game-changing, giving hope to the possibility of using bioremediation to reduce the impact of CLD contamination.

## INTRODUCTION

Chlordecone (CLD) was used to control insect pests (mainly the banana weevil) in banana plantations in the Caribbean, particularly in Guadeloupe and Martinique from 1971 to 1993, despite being banned in the United States since 1974 [1, 2]. This pesticide was also used in the USA, as well as in some countries of Africa (Cameroon, Ivory Coast), latin America (Panama, Honduras, Equator, Nicaragua) and Asia [3–5]. Chlordecone (C_10_Cl_10_O) is a bis-homocubane, comprising a cage structure with 10 chlorine atoms and a ketone functionality. It was commercialized under the brand names Kepone® and Curlone® and was spread at the foot of each banana plant as a solution delivering 1.5 grams of chlordecone per plant. With 800 banana plants per hectare and 2.5 applications per year, the resulting dosage was a significant 3 kg chlordecone per hectare per year. As a consequence of such intensive application, 8–9% of the cultivated areas of Guadeloupe have CLD concentrations higher than 1 mg/kg in topsoil, and some banana fields have concentrations higher than 9 mg/kg [2]. CLD adsorbs strongly to soils rich in organic matter, and its partitioning coefficient (log K_oc_) has been reported in the range of 3.3-3.41 [1, 6]. CLD is only slightly soluble in water (2.7 mg/L at pH 7 and 25°C [7] and tends to bioaccumulate in fatty tissues of living organisms [8, 9]. CLD is also neurotoxic, immunotoxic, hepatotoxic and spermatotoxic to most living organisms [10, 11]. In 2009, CLD was added to the list of persistent organic pollutants in the Stockholm Convention, banning its production and use worldwide. Studies suggest exposure to CLD may be linked to higher occurrence of prostate cancer and to impaired cognitive and motor development in young children [1, 12–14]. Watershed modelling predicts that CLD will remain in soils for decades or even centuries because of its stability and affinity for soil organic matter, providing a long term source of contamination to the aquatic environment [2]. Surface water, ground water, sediments and receiving coastal wasters have all been impacted. CLD has also been found to accumulate in animals [15–17] and plants used for food, including tubers and other vegetables grown in soil [18, 19] as well as in marine fauna [9, 20]. Effective methods to decontaminate soil and protect downgradient environments, food and water supply are clearly needed.

Few researchers have investigated CLD transformation. Early detailed studies by Schrauzer and Katz [21] revealed that reduced vitamin B12 could dechlorinate chlordecone into products including monohydrochlordecone (MHCLD), dihydrochlordecone (DHCLD) and pentachloroindene. Zero valent iron [22] was found to degrade CLD to different products (C_10_H_3_Cl_9_O_2_, C_10_H_4_Cl_8_O_2_, C_10_H_5_Cl_7_O_2_, C_10_H_6_Cl_6_O_2_ and C_10_H_7_Cl_5_O_2_). More recently, Ranguin et al., [23] were also able to show abiotic dechlorination using reduced vitamin B12. The abiotic reduction of CLD with B12, zero-valent iron or sodium sulphide led to formation of hydrochlordecones (HCLDs) and polychloroindenes (PCINs) [24]. Very little data exists on the microbial degradation of CLD. Using Gibbs free energy calculations, Dolfing et al. [25] showed that there is no thermodynamic reason precluding bacterial CLD degradation. Orndorff and Colwell [26] showed that *Pseudomonas aeruginosa strain K03* was able to partially transform CLD to 15% MHCLD and 5% DHCLD, however, the HCLDs detected could have been impurities and not biodegradation products [27]. Under methanogenic conditions, Jablonski et al. [28] reported extensive dechlorination and formation of apolar and polar metabolites in a culture of *Methonosarcina thermophila* at 50°C. According to this work, 86% of labelled [14C] CLD was dechlorinated. More recently, a low but significant mineralization of chlordecone under aerobic conditions was detected [29]. A fungal strain, *Fusarium oxysporum* MIAE01197, was shown to be able to grow in a liquid culture medium containing CLD as carbon source, however, no degradation products were detected [30]. Very slow, natural transformation of CLD to 5-b-hydrochlordecone was documented in polluted soils indicating the possible natural biotransformation of chlordecone [31]. More recently, the formation of MHCLD, DHCLD, trihydrochlordecone (THCLD) and indene metabolites C_9_Cl_5_H_3_ and C_9_Cl_4_H_4_ was detected in a mixed bacterial consortium as well as by an isolated *Citrobacter* strain [24, 32], and a third group of metabolites, carboxylated polycholoroindenes (CPCIN) was also reported [24]. The genome of the *Citrobacter* strain in these studies contained no reductive dehalogenase genes and thus transformation was likely cometabolic [32]. A very recent paper by the same team further identified and characterized many chlordecone transformation products [33], using chemical reduction, organic synthesis, and NMR to elucidate isomer structures. Significantly, they further detected and quantified some of these same transformation products in soil samples from Martinique.

The objective of our work was to evaluate anaerobic microbial transformation of CLD in batch bottle microcosms constructed with soil from Guadeloupe and with simulated groundwater that mimic natural attenuation and electron donor-simulated reductive dechlorination. An LC-Orbitrap MS method was eventually developed for small volume samples, where we found that the choice of extractive co-solvent was critical to the detection of both polar and non-polar metabolites. The isotopic fingerprint of compounds with multiple chlorines enabled identification of metabolites. Microcosms were maintained under strictly anaerobic conditions, and we were able to document slow but extensive dechlorination of CLD to PCIN and CPCIN metabolites with up to 9 chlorine atoms removed. A crude estimation of partitioning coefficients between soil and water was possible to provide first insights into the fate and transport of these CLD metabolites. With a suitable detection method in hand, we then analyzed several field samples from Guadeloupe. In these samples, MHCLD, DHCLD and pentachloroindene metabolites were convincingly detected, indicating that these dechlorinating reactions can also proceed *in situ*, most likely where anaerobic field conditions can be found.

## MATERIALS AND METHODS

### Collection of Field Samples

Field samples were collected twice, first in the fall of 2010 for microcosm setup, and second in the spring of 2018 to analyze CLD and potential degradation products by LC/MS using a refined protocol. In 2010, samples were collected from seven locations in the south of Basse-Terre island, Guadeloupe (Table S1A): three andosol samples from agricultural areas used for banana production (soil); three fluvisol samples from river banks near a banana production area (soil and water); and one sludge sample from an anaerobic digester at a sugar cane distillery plant. In 2018, nine different locations were sampled (Table S1B): six from CLD-impacted agricultural areas (as soil and water slurries); and three activated carbon sludge samples from a water treatment plant that handles chlordecone-contaminated water (sampling details in Supplemental Method Details (SMD) 1A).

### Chemicals

CLD (neat) and CLD standard (analytic standard 1mg/mL in MeOH) were purchased from Accustandard (New Haven, USA). Hexanes and acetone (Fisher), water and methanol (Caledon Laboratory Chemicals), and ethanol (Commercial Alcohols, Brampton, Canada) were all of HPLC grade. TCE was purchased from Sigma and had a purity of 99.5%. A pentachloroindene standard (2,4,5,6,7-pentachloro-1H-indene) referred to as B1, was kindly provided by researchers at Genoscope (Évry, France).

### Microcosm Setup and Enrichment

In December 2010, a microcosm study consisting of 13 different conditions each in triplicate for a total of 39 microcosms was initiated (Table S2A). Microcosms contained soil and water samples from Guadeloupe and were augmented with artificial groundwater (recipe in SMD 1B) as there was not enough field water. One set of microcosms contained sludge from an anaerobic digester instead of soil. The microcosms were prepared in 160 ml serum glass bottles (Fisher Scientific) sealed with blue butyl stoppers (Bellco Glass Inc.) with 22.5 ml soil and 80 ml water. Rezasurin (1 mg/l) was added to one microcosm from each triplicate set as an indicator of anaerobic conditions. At setup, 15 microcosms (4 different soils and one sludge sample, all in triplicate) were poisoned by adding mercuric chloride (0.05%) and sodium azide (0.02%). The 15 poisoned control microcosms and 18 of the 24 active microcosms were amended with CLD at a target concentration of 10 mg/l (which is above the solubility of ~2 mg/L [1, 8], therefore a separate phase of CLD was expected). Three other active microcosms were amended with both CLD and trichloroethene (TCE) at target concentrations of 10 mg/l each, and three microcosms were left unamended. TCE was added as a positive control for reductive dechlorination. All microcosms, except from the three unamended bottles, received an initial dose of an electron donor mix of acetone and ethanol, each at ~80 mg/L. The ratio of donor (acetone/ethanol) to CLD (acceptor) in terms of electron equivalents (eeq) was around 100:1, assuming 20 eeq/mol CLD for complete dechlorination. This ratio is very high, and higher than typically used for other chlorinated electron acceptors like TCE (5:1) because more acetone and ethanol were needed to dissolve CLD (solid powder) into the feeding stock solution (Table S3). The microcosms were incubated in the dark, unshaken in an anaerobic glovebox (Coy Lab Products, Grass Lake, MI, USA) for about 8 years. During the first 2.5 years, all the active microcosms were re-amended with CLD twice and electron donor five times. Those microcosms that also received TCE were re-amended with TCE twice during this period. Six active microcosms were transferred into a defined pre-reduced anaerobic mineral medium [34] in slightly larger bottles: one after 1 year, one after 1.5 years, and four after 2.5 years (Figure S1 and Table S2B). These transfers were made in 250 ml Boston Round glass bottles (VWR) to a total liquid volume of 200 ml and sealed with screw-cap Mininert™ valves (Chromatographic specialties). Subsequently, these transferred bottles (GT5, GT20, GT33, GT4, GT15, GT3) were re-amended with donor and acceptor regularly. All the other microcosms were not maintained after 2.5 years, but some were sampled a few times for comparison with the transfers. The amounts and frequency of amendments to original microcosms and to transfers over the full study are shown in Table S4. Two bottles with medium only (Medium1, Medium2) were prepared in 2016 for use as controls for LC/MS analysis. These were set up in 250 ml Boston Round glass bottles with 200 ml mineral medium (no culture) and amended with CLD feeding stock to CLD concentrations of 10 and 20 mg/l (Table S2C).

### Microcosm Sampling and Analysis

For the first 1.5 years, liquid samples (1 ml) from microcosms were sampled regularly and analyzed for methane, ethene, ethane, and chlorinated ethenes by Gas Chromatography with Flame Ionization Detector (GC-FID), but beyond this time, only the transferred microcosms were analyzed routinely.

Anion analysis (IC) and pH measurements were made a few times over the course of the study. Samples for DNA extraction were taken from the 6 transfers (GT5, GT20, GT33, GT4, GT15, GT3) about 7 years into the study and the microbial community composition was assessed by small subunit (SSU) rRNA gene amplicon sequencing and Quantitative Polymerase Chain Reaction (qPCR) analysis. Method details of GC, IC, pH and microbial community analysis are described in SMD 2.

When the microcosms were first established, we did not have a good method to measure CLD or its daughter products. Nevertheless, during the first 1.5 years, 1 ml liquid samples (in duplicate) were taken every 1-2 months from each microcosm and archived frozen at −20°C. Once an appropriate LC/MS method was developed, analyses by LC/MS were performed approximately once per year, and more frequently in the last two years. The transfers (GT5, GT20, GT33, GT4, GT15, GT3) were analyzed most often, and some poisoned controls, medium controls and some of the original active microcosms were analyzed occasionally for comparison. Sampling and sample preparation procedures for the LC/MS analysis are described below with further details in SMD 2.

### Sampling and Sample Preparation for CLD Analysis by LC/MS

Sampling procedures, sample preparation methods and chromatographic and MS methods for analysis of CLD and dechlorinated products were improved progressively (explained in SMD 2). Eventually, two satisfactory sample preparation methods were developed for the microcosms, one for sampling liquid only (method 3) and one for sampling a soil/water slurry (method 4) (details in SMD 2). Samples were always taken using glass syringes (Hamilton Company, Reno, USA) and stored in glass vials with caps with PTFE lined septa (Agilent) to minimize sorption. All samples and standards were filtered through 0.2 μm Millex PTFE syringe filters (Millipore, Burlington, USA). For the liquid samples (method 3), bottles were shaken and left to settle overnight. The next day 0.75 ml liquid was carefully sampled (avoiding collecting any solids) and was placed into a 2 ml glass vial. The sample was then centrifuged for 5 minutes at 3000 rpm, and 0.5 ml of the supernatant was transferred into a new vial containing 0.5 ml methanol. After mixing, the sample was filtered into a final 2 ml glass sampling vial. For the slurry samples (method 4), bottles were shaken and a 1 ml sample was taken with syringe immediately, before the solids settled to get a representative sample of the soil/water slurry. The slurry sample was added to a vial containing 1 ml methanol, shaken gently for 10 min and left sitting for 30 min. The sample was then centrifuged for 5 minutes at 3000 rpm, and the supernatant was filtered into the final 2 ml sampling vial. The addition of methanol to the sample had three purposes; to help extract compounds associated with the solids, to help clean up the sample matrix by inducing precipitation of salts from the sample matrix, and to reduce sorption of compounds to filter membranes. For some of the samples the dry weight of solids was measured after drying the sample in a drying oven at 105°C for several hours to estimate sorbed mass.

The field samples taken in 2018 were expected to have much lower concentrations of CLD and daughter products than the microcosms, therefore a different extraction and concentration procedure was used. Two different sample volumes (5 ml and 20 ml) were extracted by liquid-liquid extraction using a mix of 15% acetone and 85% hexane (methods 5 & 6). The extracts were evaporated to dryness under a stream of nitrogen and the samples were re-dissolved in 1ml and 0.5ml methanol and filtered. Sample preparation procedures, LC/MS instrumentation and methods were in constant development over the course of the study. Only the final optimized method, used from April 9^th^ 2018, is described in detail here. The history and development of the protocol is explained in SMD 2.

### Liquid Chromatography Mass Spectrometry (LC/MS) Analysis

Chromatography was carried out on a ZORBAX Rapid Resolution High Definition Phenyl-Hexyl column (50mm x 3.0mm, 1.8um) (Agilent, Santa Clara, USA) equipped with a guard column, using a Thermo Scientific Ultimate 3000 UHPLC (Thermo Fisher Scientific, Waltham, MA). The column temperature was 40°C and the flow rate was set to 300 μL·min^−1^. The eluents used were water (A) and methanol (B), and both eluents contained 5 mM of ammonium acetate. The gradient started at 50% B, followed by a linear gradient to 100% B over 8 min, then a hold at 100% B for 4 min, then a return to 50% B over 1min, and finally a re-equilibration under the initial conditions of 50% B for 5 min (total runtime 18 min). Liquid samples (10 μL) were injected using an Ultimate 3000 UHPLC autosampler, with autosampler temperature of 8°C. Compounds were detected and quantified using a Q-Exactive Orbitrap mass spectrometer (Thermo Fisher Scientific) equipped with a Heated Electrospray Ionization (HESI II) probe, operating in negative ionization mode. Mass spectra were acquired over an m/z range from 80 to 750 with the mass resolution set to 140k, and common setting parameters were as follows; AGC Target: 1E6, max injection time 200 ms, spray voltage 4 kV, capillary temperature 320°C, sheath gas 15, aux gas 5, spare gas 2, and s-lens RF level 55. Data from the LC/MS were processed through Xcalibur Qual Browser (Thermo Fisher Scientific). Calibration standard solutions of CLD (0.02, 0.2, 1.7, 2.6, 3.6 mg/l) were prepared from successive dilutions of a purchased analytical standard (1 mg/mL CLD in MeOH). Similarly, a sample of pentachloroindene B1 (2,4,5,6,7-pentachloro-1H-indene) (from Genoscope, France), stored as a 0.4 mg/ml acetonitrile stock solution, was diluted to make five concentration level (0.02, 0.1, 0.2, 0.5 and 1 mg/l) and used as standards for estimates of CLD metabolite concentrations. The dilutions of CLD and B1 were prepared in a 50/50% mix of methanol and mineral medium. Standard dilutions were made fresh the day before or on the same day as they were run.

## RESULTS AND DISCUSSION

### Overview of Microcosm Data

CLD is a challenging molecule to quantify, particularly in small volume samples from a microcosm. It took us several years after microcosm set up to develop a suitable method, and until that time, we periodically amended microcosms with CLD without knowing if it was being degraded or not. We rationalized that dechlorination would be slow and limited by the low solubility of CLD. Therefore, provided that we had good controls, we would eventually be able to identify activity by comparing active microcosms to poisoned and medium controls.

To provide an overview of how each set of microcosms was treated over the course of about 3000 days (~8 years), cumulative CLD and electron donor (acetone and ethanol) amendments were plotted over time (Figure S2; with data in Table S4). We monitored methane production, sulfate consumption and TCE dechlorination (where added), as indications of anaerobic microbial activity. All active microcosms except for those made from distillery sludge samples (G63 series) produced methane during the first 1.5 years (Figure S3). Beyond 1.5 years, we continued to monitor and amend only six of the microcosms. Two (GT4 and GT3) were 20% transfers from original bottles into anaerobic medium, that received CLD and donor (GT4) or CLD+TCE and donor (GT3). Transfer GT4 contained none of the original microcosm soil. Four other transfers were made by pouring the entire contents of the original microcosm into slightly larger bottles sealed with Teflon Mininert caps, topped up with medium and amended with CLD and donor (GT5, GT20 and GT33) or CLD+TCE and donor (GT15) (Figure S1). Transfers GT5, GT20 and GT15 continued to produce methane after being transferred, GT33 stopped producing methane after the first re-amendment of CLD, while GT3 and GT4 did not produce any methane after the transfer (Figure S2). In microcosms that received TCE in addition to CLD, TCE was partially dechlorinated to trans- and cis-DCE, with possibly small amounts of ethane produced (Figure S4), but the 20% transfer (GT3) no longer dechlorinated TCE. No ethene was ever found in any of the microcosms (Table S5). In the transfers that did not produce methane (GT33, GT3 and GT4), we observed production of acetate (Table S5) although not enough to explain complete transformation of donor. As a result, pH dropped to below 6 in these microcosms (and was occasionally re-adjusted), which probably subsequently eliminated methanogens (Table S5). Sulfate (~1.2 mM initially) was present in all soil microcosms but was depleted within the first year and was not added to transfers (Table S5). We suspect that iron(III) carried over from soil may have also served as an acceptor in some bottles. High chloride background (>12 mM) was present in the microcosms, precluding using chloride increase as a proxy for dechlorination. In summary, we established that all of the active (non-poisoned) microcosms were microbially active initially as well as after 8 years of incubation, despite repeated additions of chlordecone and donor (acetone or ethanol), but that there were significant differences in their extent of methanogenesis depending on particular treatment history.

### Development of an LC/MS Method for Chlordecone and metabolites

To enable a time-course analysis and mass balance in microcosms, we needed to develop a CLD analysis method suitable for repeated sampling in microcosms using small sample volumes. CLD analyses are challenging for many reasons, including strong sorption of CLD to soil (and to rubber stoppers), low water solubility of CLD and certain metabolites, lack of authentic standards for degradation products, poor/variable or inconsistent ionization, as well as matrix and solvent effects. We hoped that the high sensitivity and mass accuracy of Orbitrap MS technology would facilitate identification of compounds despite lack of standards. The various early trials and final method are described in SMD 2. Moriwaki and Hasegawa [35] were first to report use of LC/MS for CLD; they used a water/methanol gradient and negative ionization mode, although only standards in methanol were tested. Durimel *et al.,* [36] also detected CLD by LC/MS using a water/acetonitrile gradient, also in negative mode. Cimetiere et al., [37] included formic acid (pH 2) to the eluent and observed adduct (M+HCOOH+OH) formation. These authors recommended addition of a co-solvent (acetonitrile, methanol) to avoid adsorption of CLD to filters and to desorb CLD from suspended matter (acetonitrile). They concluded that the presence of salts in sample matrixes weaken the CLD MS signal, which made us realize the importance of preparing standards in the same matrix as the samples.

Based on these prior studies, standards were prepared in 50% medium and 50% methanol. Methanol was added to samples before filtration to decrease sorption and to precipitate salts. Methanol addition proved later to be a good solvent to mix with water to recover compounds from soil particles. As first described by Harless et al., [38] and observed by others [24, 39], CLD actually exists as a gem-diol (hydrate) in water or a hemiacetal in methanol and can also form various adducts in the presence of compounds often used in LC eluents, such as acetate and formate. In this study chromatographic separation, ionization and signal intensity were maximized when ammonium acetate was included in the water and methanol eluents. We observed many CLD adducts (hydrate, formate, acetate) and hemiacetal in our analyses (Figure S5), that required careful data interpretation. Hydrochlordecones (MHCLD, DHCLD and THCLD) also showed similar adduct formation. In this study we chose to calibrate and report the hydrate forms of CLD, MHCLD, DHCLD and THCLD. All of the LC/MS data collected during this study are compiled in Table S7. We initially only sampled the liquid phase (centrifuged and filtered) to avoid variability related to sorption to solids in the microcosms. However, sorption to soil in the microcosm was very strong, precluding a satisfactory mass balance when considering only liquid phase samples. We eventually had to adapt the method to include and quantify CLD on solids. Despite all these challenges, dechlorination products were clearly observed, as described in the next section.

### Detection of CLD Metabolites using best method

A series of metabolites were observed in all the active original microcosms and transfers over the course of the study (Figure 1 with data in Table S7). Despite not having standards, the accuracy of the Orbitrap MS enabled identification of metabolites by matching first or highest m/z values of the mass spectrum to the presumed structure, and further evaluating the full characteristic isotope patterns for these multi-chlorinated transformation products (details below). Table S6 lists all observed metabolites with their measured and theoretical masses (and IDs), and Figure S6 illustrates the characteristic mass spectra for these metabolites (observed and theoretical). The identified metabolites were assigned to three different groups: group A - hydrochlordecones (HCLD); group B - polychloroindenes (PCIN); and group C - carboxylated polychloroindenes (CPCIN) to be consistent with previous studies detailing the masses and NMR structures of CLD metabolites [24, 32, 40]. In this study, a total of 19 different dechlorination products were detected by LC/MS in the microcosms. Multiple isomers of certain metabolites were found, having exactly the same mass but varying retention time. Retention times were also consistent with the relative polarity of each compound. In this study we relied on exact masses, unique isotope distribution patterns, and retention times to support the indentity of metabolites classes, namely hydrochlordecones, polychloroindenes, and carboxylated polychloroindenes with varying numbers of chlorine substituents. The consistency of the results with previous reports [24, 33] and highly characteristic chlorine isotope pattern leave little doubt to the structure of the compounds, other than the actual position of substituents on the rings. Only purification of each compound and NMR could resolve such structural details, and these are beyond the scope of this work, and not necessary to evaluate extent of dechlorination. An example chromatogram from one of the samples, bottle GT20 sampled June 29^th^ 2018, is shown in Figure 2. Chlordecone and 17 of the 19 observed dechlorination products detected in this study are shown, including the non-polar B compounds (polychloroindenes, PCINs) showing longer retention times than CLD, and the more polar C compounds (carboxylated polychloroindenes, CPCINs) with much shorter retention times (Figure 2). Compounds with retention times 9.3, 8.88, and 8.05 min and mass to charge ratios m/z 468.7264, m/z 434.7661 and m/z 400.8042 respectively, could be attributed to the hydrate forms of MHCLD (A9a) [C_10_Cl_9_O_2_H_2_]^−^, DHCLD (A8a) [C_10_Cl_8_O_2_H_3_]^−^ and THCLD (A7a) [C_10_Cl_7_O_2_H_4_]^−^ obtained by the loss of 1, 2 and 3 chlorine atoms from CLD (Figure 2 and Table S6). Three metabolites, less polar than CLD and with longer retention times at 12.19, 11.44 and 10.34 min, respectively, exhibiting mass to charge ratios of m/z 284.8616, m/z 250.9006 and m/z 216.9382 were identified as pentachloroindene (B5a) [C_9_Cl_5_H_2_]^−^, tetrachloroindene (B4a) [C_9_Cl_4_H_3_]^−^ and trichloroindene (B3a) [C_9_Cl_3_H_4_]^−^ (Figure 2 and Table S6). Thirteen (13) metabolites more polar that CLD were assigned to group C, and observed at retention times ranging from 2.02 to 6.69 min (Figure 2 and Table S6). Compounds C4a and C4b, detected respectively at retention times of 5.47 and 5.02 min, and with a corresponding m/z of 294.8899 could be attributed to two isomers of a carboxylated tetrachloroindene compound [C_10_Cl_4_O_2_H_3_]^−^ obtained by the loss of 6 chlorine atoms. At retention times between 3.03 and 6.69 min, 5 isomers (C3a-C3e) with a corresponding m/z of 260.9287 were detected and attributed to a carboxylated trichloroindene compound [C_10_Cl_3_O_2_H_4_]^−^ obtained by the loss of 7 chlorine atoms. Four isomers (C2a-C2d) obtained by the loss of 8 chlorine atoms were detected at retention times between 2.02 and 5.25 min and had m/z of 226.9677, assigned to be a carboxylated dichloroindene [C_10_Cl_2_O_2_H_5_]^−^. Two last isomers (C1a and C1b) with retention times at 2.5 and 2.95 min and m/z of 193.0064 were classified as carboxylated monochloroindene [C_10_ClO_2_H_6_]^−^, obtained by the loss of 9 chlorine atoms from CLD.

**Figure 1:**
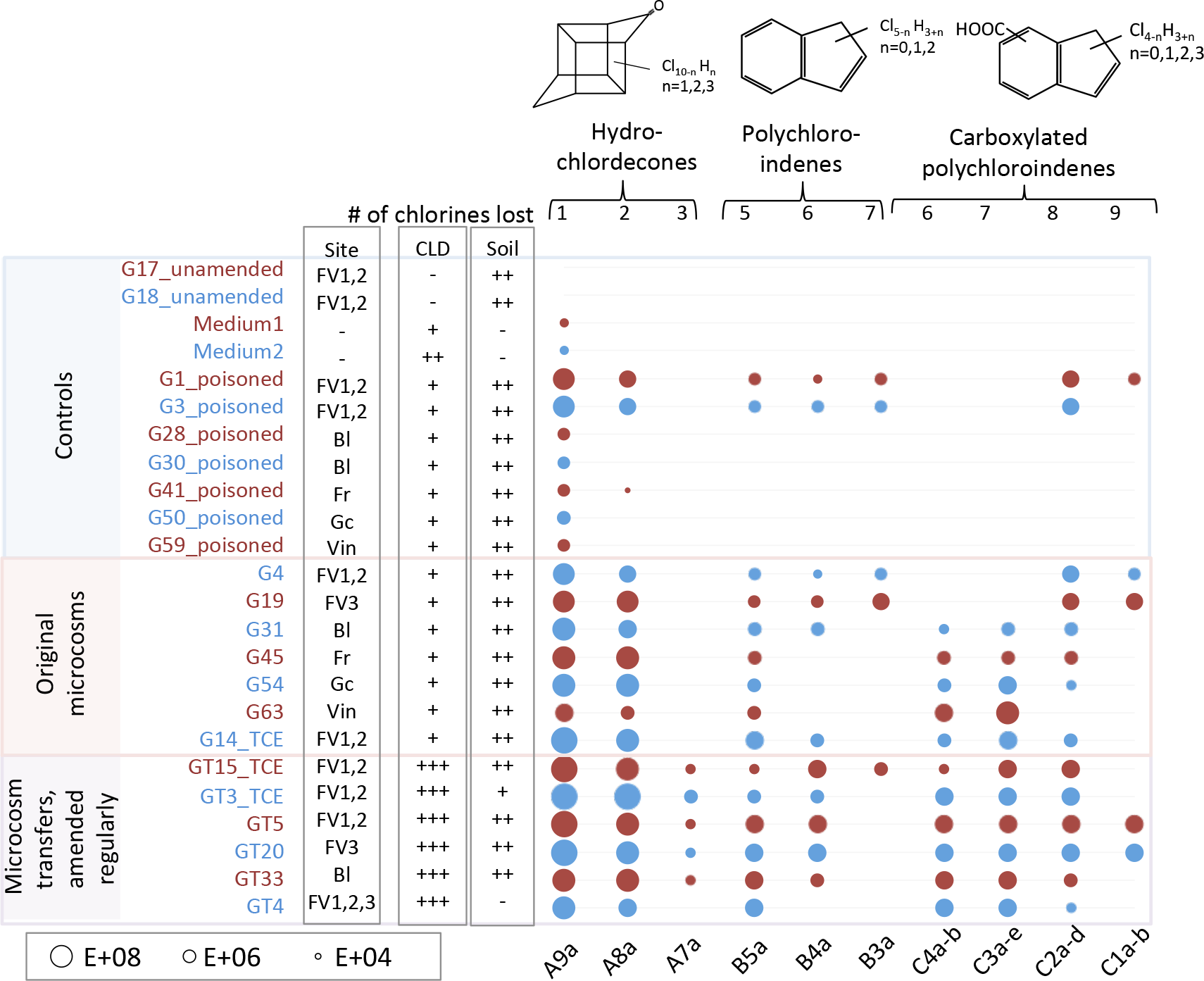
Dechlorinated metabolites observed in anaerobic microcosms constructed from Guadeloupe soil and water. The size of each circle is proportional to the area count from LC/MS analysis of slurry samples performed March 7^th^ 2019 (day 3038) (see Table S7 for raw data).

**Figure 2:**
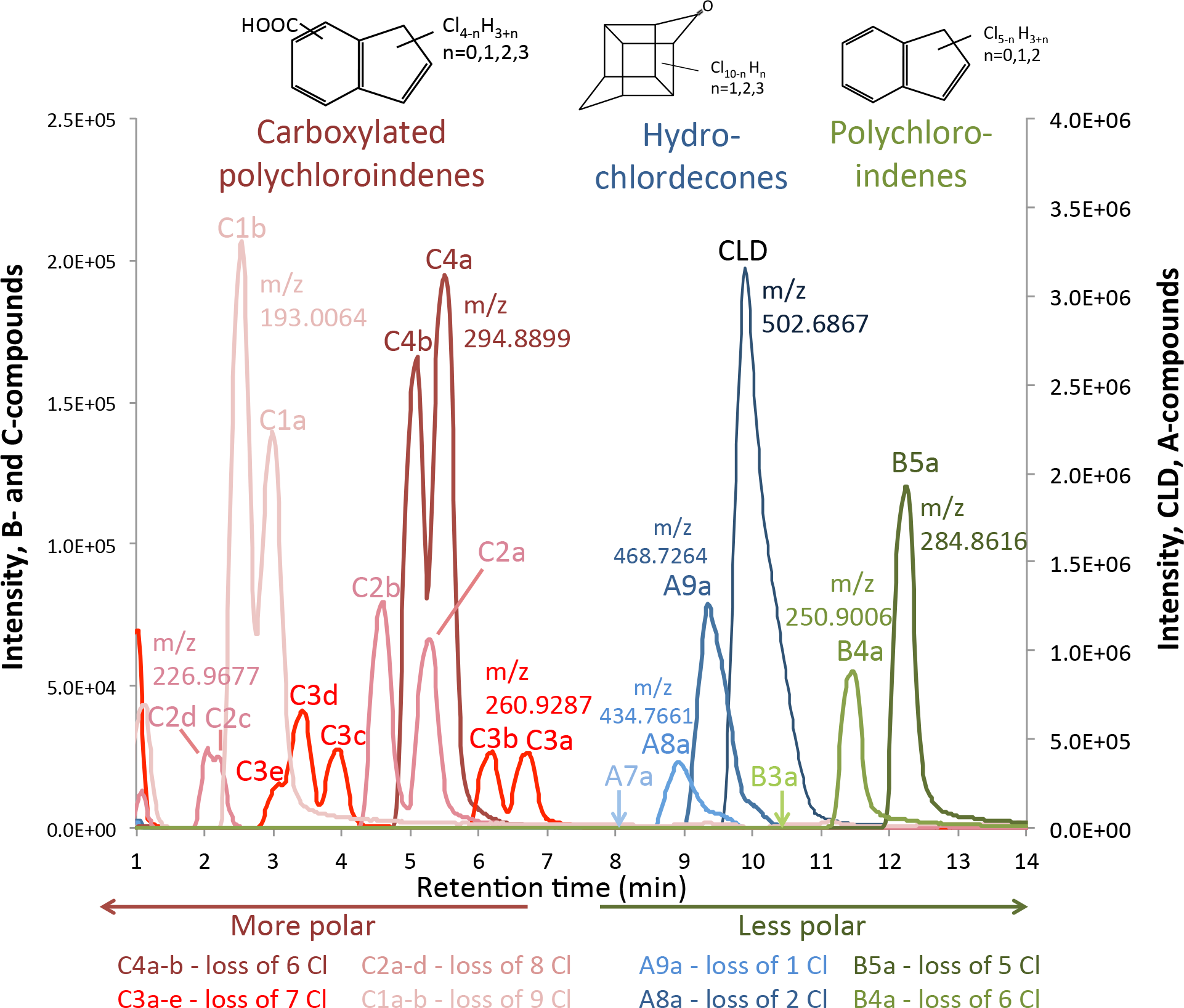
Chromatograms of chlordecone and its dechlorinated metabolites in sample GT20 June 29^th^ 2018 (slurry sample). Results are from LC/MS, equipped with ESI, in negative mode. Observed m/z values (monoisotopic) for the different compounds were; 502.6879 (CLD, [C_10_Cl_10_O_2_H]^−^), 468.7264 (A9a, [C_10_Cl_9_O_2_H_2_]^−^), 434.7661 (A8a, [C_10_Cl_8_O_2_H_3_]^−^), 284.8616 (B5a, [C_9_Cl_5_H_2_]^−^), 250.9006 (B4a, [C_9_Cl_4_H_3_]^−^), 294.8899 (C4a-b, [C_10_Cl_4_O_2_H_3_]^−^), 260.9287 (C3a-e, [C_10_Cl_3_O_2_H_4_]^−^), 226.9677 (C2a-d, [C_10_Cl_2_O_2_H_5_]^−^), 193.0064 (C1a-b, [C_10_ClO_2_H_6_]^-^) (see details in Table S6). Metabolites m/z 400.8042 (A7a, [C_10_Cl_7_O_2_H_4_]^−^) and 216.9382 (B3a, [C_10_Cl_7_O_2_H_4_]^−^) were not observed in the illustrated sample but in some other samples in the study. Arrows indicate observed retention times for these two metabolites. CLD, MHCLD, DHCLD and THCLD are quantified/reported in the forms of CLD hydrate, MHCLD hydrate, DHCLD hydrate and THCLD hydrate.

In summary, CLD metabolites observed in the microcosms could be classified into 3 families of compounds: group A compounds including mono-, di- and trihydrochlordecone derivatives with proposed neutral formula [C_*10*_Cl_*10*−*n*_OH_*n*_, n= 1,2,3]; group B non-polar “open cage” structures including three polychloroindene compounds with proposed neutral formula [C_*9*_Cl_*5*−*n*_H_*3+n*_, n=0,1,2]; and group C polar “open cage” structures consisting of carboxylated polychloroindene derivatives with neutral formula [C_*10*_Cl_*4*−*n*_O_*2*_H_*4*+*n*_, n=0−3].

Due to a number of challenges with and changes in sample preparation and LC/MS analysis over time, it was difficult to get an accurate picture of concentration changes of CLD over time. Therefore, when we finally settled on a good method, we re-analyzed some frozen archived samples to get a comparable set of data. Results from a selection of samples from GT20 with comparable sample preparation (liquid phase only, no soil) and analytical methods are shown in Figure 3 (raw data in Table S8). Results show that none of the monitored metabolites were detected in the sample taken two weeks into the experiment, but by 8 months into the study, we observed MHCLD, DHCLD, and two CPCINs with loss of 8 and 9 chlorines. Later sampling time points showed increasing concentrations of MHCLD, DHCLD, and four CPCINs with loss of 6 to 9 chlorines. Pentachloroindene (B5a) only showed up in the last sampling point, however it was found to sorb strongly to soil, and we would therefore not expect to see it in these liquid phase samples analyzed here. Also, because we kept adding more CLD to the bottles over time, we could not use aqueous CLD concentration changes as a measure of degradation rate. Despite not seeing a clear decrease in CLD concentrations in the liquid phase, highly dechlorinated metabolites with up to 9 chlorine removed were observed in the active bottles but not in the controls indicating that biological processes were involved in the dechlorination of CLD into HCLD, PCIN and CPCIN metabolites.

**Figure 3:**
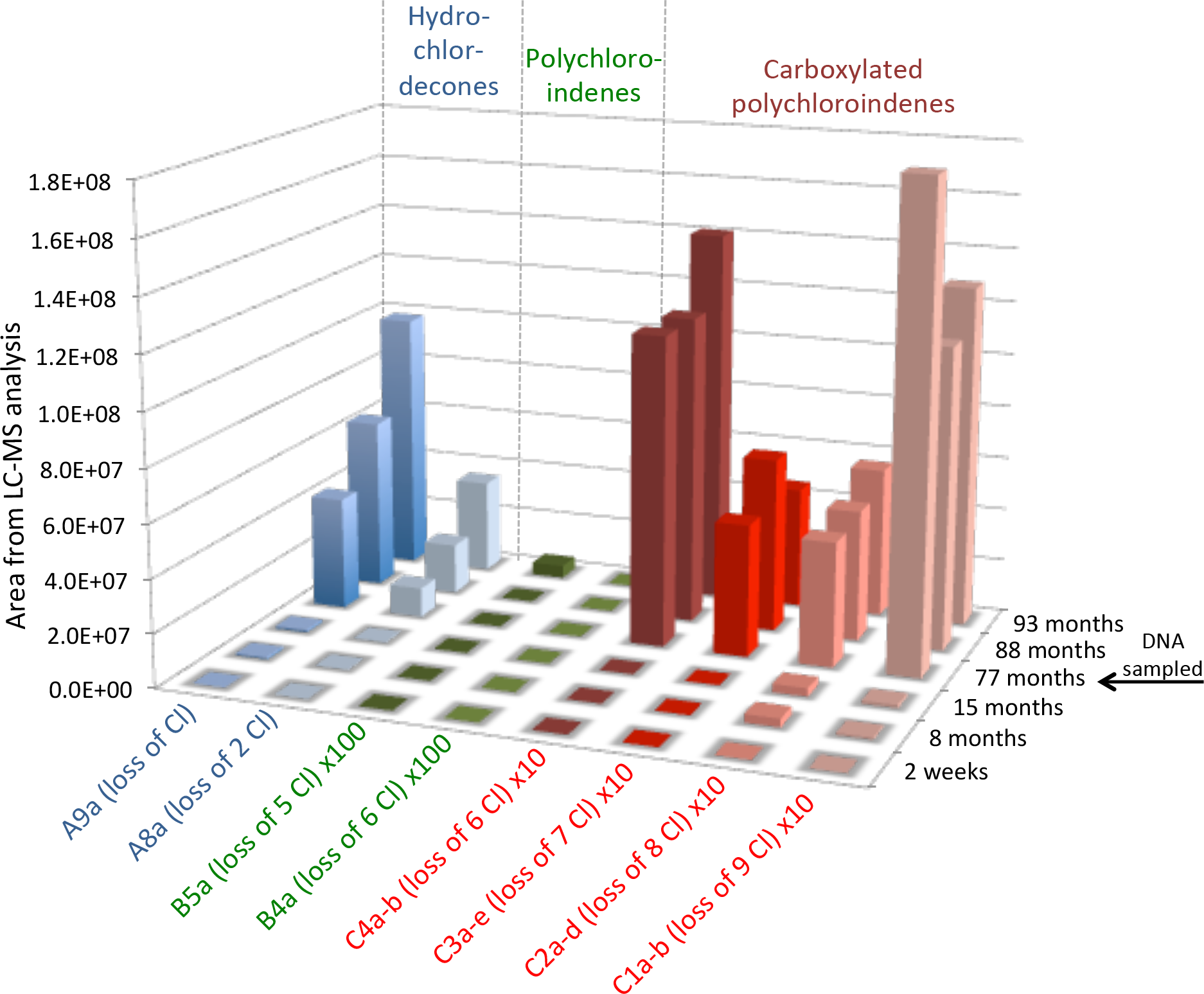
Dechlorinated metabolites from chlordecone over time in one of the anaerobic microcosms, transfer GT20. Only samples that were prepared the same way are included in this graph (sample preparation method 3, liquid phase only, no soil, see supplemental method details). Areas of B compounds were multiplied with 100, and areas of C compounds were multiplied with 10 for better display of all metabolites in the same graph. All areas were normalized (raw data in Table S8).

### Effect of Sorption

Strong sorption of CLD and non-polar PCID metabolites to soil particles made it difficult to evaluate the fate of CLD in the bottles by sampling the aqueous phase only. We therefore changed our approach to also extract soil in our samples. By analyzing the same samples by two different sample preparation methods, one in which only the liquid concentration was measured (filtered sample) and another in which a mixture of soil and water was analyzed, and quantifying the amount of soil (dry mass) in the sample, we were able to calculate and estimate a distribution coefficient, K_d_ (l/kg) equal to the ratio of sorbed concentration (mg/kg) to dissolved concentration (mg/l) for most of the analytes (Table 1; raw data in Table S9). This analysis confirmed that CLD and mono- and dihydro- CLD, with K_d_ values of 130 ± 57, 52 ± 12 and 28 ± 6 ml/g (or l/kg), respectively, absorb quite strongly to the Guadeloupe soil used, while the K_d_ value for pentachloroindene (B5a) was much higher at 5700 ± 220 l/kg, and thus absorb even more strongly. The carboxylated chloroindenes (C compounds) had much lower K_d_ values ranging from 2 to 11 l/kg, and were found in the aqueous phase. The estimated K_d_ values correspond well to the retention times by reverse phase LC (Table 1). Our estimated distribution coefficient for chlordecone is in the same range (60-330 l/kg) as one previous report [41]. A distribution coefficient based on organic content, K_oc_, of 2,500 l/kg has also been reported [8], corresponding to a K_d_ of 250 l/kg assuming a fraction of organic carbon of 10%. We were not able to find any reports of sorption coefficients for any CLD metabolites.

**Table 1:**
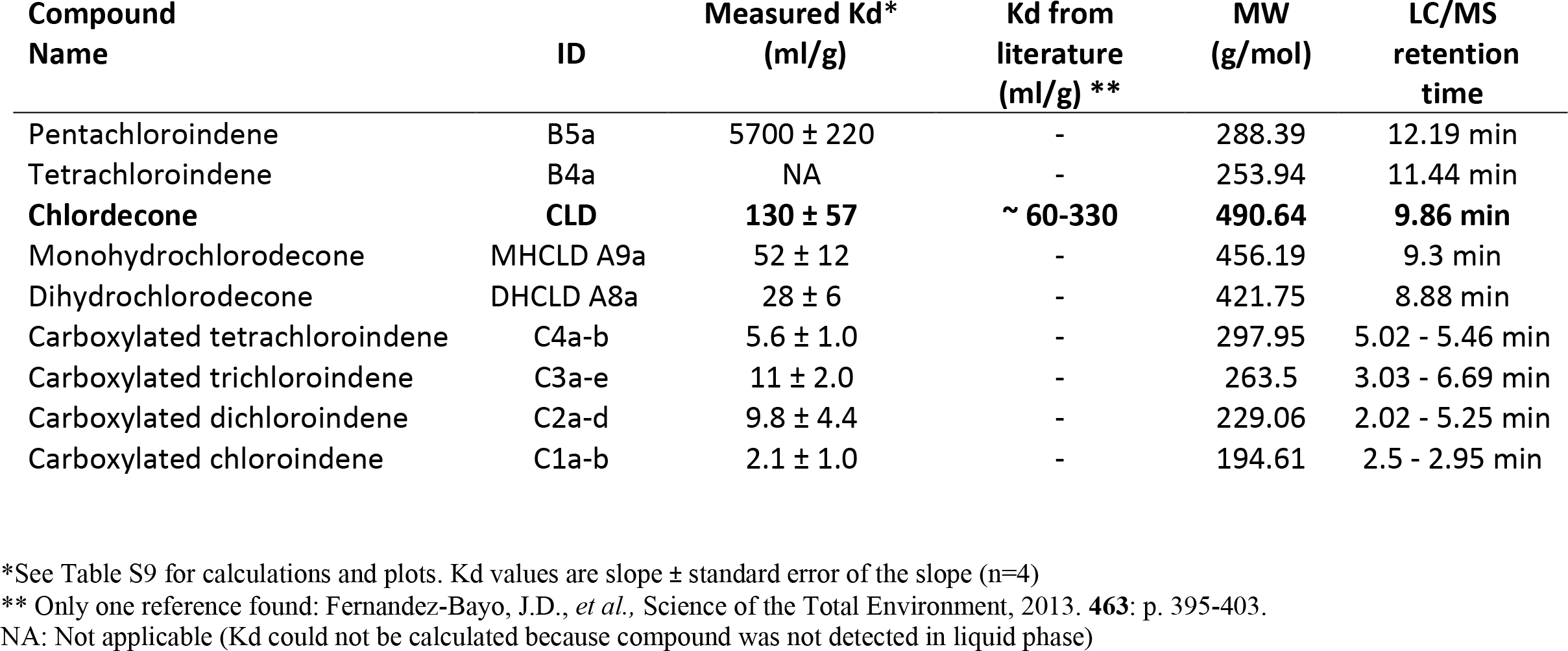
Estimated sorption coefficients for CLD and some dechlorinated metabolites in Guadeloupe soils

| Compound Name | ID | Measured Kd*<br>(ml/g) | Kd from literature<br>(ml/g) ** | MW<br>(g/mol) | LC/MS retention time |
| --- | --- | --- | --- | --- | --- |
| Pentachloroindene | B5a | 5700 ± 220 | - | 288.39 | 12.19 min |
| Tetrachloroindene | B4a | NA | - | 253.94 | 11.44 min |
| **Chlordecone** | **CLD** | **130 ± 57** | **~ 60-330** | **490.64** | **9.86 min** |
| Monohydrochlorodecone | MHCLD A9a | 52 ± 12 | - | 456.19 | 9.3 min |
| Dihydrochlorodecone | DHCLD A8a | 28 ± 6 | - | 421.75 | 8.88 min |
| Carboxylated tetrachloroindene | C4a-b | 5.6 ± 1.0 | - | 297.95 | 5.02 - 5.46 min |
| Carboxylated trichloroindene | C3a-e | 11 ± 2.0 | - | 263.5 | 3.03 - 6.69 min |
| Carboxylated dichloroindene | C2a-d | 9.8 ± 4.4 | - | 229.06 | 2.02 - 5.25 min |
| Carboxylated chloroindene | C1a-b | 2.1 ± 1.0 | - | 194.61 | 2.5 - 2.95 min |

\*See Table S9 for calculations and plots. Kd values are slope ± standard error of the slope (n=4)
\*\* Only one reference found: Fernandez-Bayo, J.D., *et al.*, Science of the Total Environment, 2013. **463**: p. 395-403.
NA: Not applicable (Kd could not be calculated because compound was not detected in liquid phase)

### Quantification of the Extent of Transformation in Microcosms and mass balance calculations

To quantify the extent of transformation of CLD added to the microcosms over the course of the study, we used data from well-mixed slurry samples from 13 active, 7 poisoned controls, 2 unamended microcosms, and two medium controls (Table 2). The well-mixed slurry samples were analyzed because they capture mass from both the liquid and solid phases, so that we could better compare final measured mass recovered to the total amount of CLD that had been added to the bottles. To attempt a mass balance, MHCLD, DHCLD, and THCLD concentrations were estimated based on the response factor for CLD as no standards were available for these metabolites. Researchers from Genoscope (France) kindly provided us with a small sample of pentachloroindene B1 (2,4,5,6,7-pentachloro-1H-indene) that they had managed to chemically purify. As a result, we were able to do a rough estimate of the amount of B5a produced in active microcosms. B4a concentrations were estimated using the response factor for B1. For the carboxylated polychlorinated indenes (C group), we have no proxy for calibration; however, to get a very rough idea of possible concentrations we also used the response factor for B1 for these compounds.

**Table 2:** Extent of transformation of CLD in microcosms and transfers (μmol per bottle) after 8 years of incubation. Concentrations of metabolites are estimates. Samples were taken March 7th 2019 (1 ml slurry samples; sample preparation method 4). Raw data and calculations are shown in Table S10.

|  | Sample | Total CLD added to date | CLD remaining after 8 yrs | CLD recovery | MHCLD, DHCLD and THCLD* (Estimated) | PCINs (B-comp)** (Estimated) | CPCINs (C-comp)** (Estimated) | Sum of all products (Estimated) | Sum all after 8 years (incl. CLD) (Estimated) | Products/CLD remaining (Estimated) | Total recovery after 8 years (Estimated) |
| --- | --- | --- | --- | --- | --- | --- | --- | --- | --- | --- | --- |
|  |  | (µmol) | (µmol) | % | (µmol) | (µmol) | (µmol) | (µmol) | (µmol) | % | % |
| CONTROLS | G17_unamended | 0.00 | 0.00 | NA | ND | ND | ND | 0.0 | 0.00 | 0% | NA |
|  | G18_unamended | 0.00 | 0.00 | NA | ND | ND | ND | 0.0 | 0.00 | 0% | NA |
|  | Medium1 | 4.08 | 4.0 | 98% | ND | ND | ND | 0.0 | 4.0 | 0% | 98% |
|  | Medium2 | 8.15 | 1.7 | 21% | ND | ND | ND | 0.0 | 1.7 | 0% | 21% |
|  | G28_poisoned | 1.63 | 1.1 | 69% | 0.002 | ND | ND | 0.002 | 1.1 | 0% | 69% |
|  | G30_poisoned | 1.63 | 0.96 | 59% | 0.001 | ND | ND | 0.001 | 0.97 | 0% | 59% |
|  | G41_poisoned | 1.63 | 1.1 | 65% | 0.004 | ND | ND | 0.004 | 1.1 | 0% | 65% |
|  | G50_poisoned | 1.63 | 0.90 | 55% | 0.002 | ND | ND | 0.002 | 0.90 | 0% | 55% |
|  | G59_poisoned | 1.63 | 1.1 | 65% | 0.001 | ND | ND | 0.001 | 1.1 | 0% | 65% |
| **AVERAGE poisoned contr.** |  | **1.63** | **1.0** | **63%** | **0.002** | **ND** | **ND** | **0.002** | **1.0** | **0.2%** | **63%** |
| **Standard Dev (n-5)** |  | **0.00** | **0.09** | **6%** | **0.001** | **-** | **-** | **0.001** | **0.09** | **0.1%** | **6%** |
| ORIGINAL MICROCOSMS | G1_poisoned# | 1.63 | 0.83 | 51% | 0.092 | 0.030 | 0.10 | 0.22 | 1.0 | 27% | 64% |
|  | G3_poisoned# | 1.63 | 0.83 | 51% | 0.079 | 0.061 | 0.08 | 0.22 | 1.1 | 26% | 64% |
|  | G4 | 3.26 | 1.4 | 42% | 0.14 | 0.047 | 0.11 | 0.30 | 1.7 | 22% | 51% |
|  | G19 | 3.26 | 1.7 | 53% | 0.45 | 0.13 | 0.24 | 0.81 | 2.6 | 47% | 78% |
|  | G31 | 3.26 | 1.7 | 53% | 0.13 | 0.045 | 0.05 | 0.22 | 1.9 | 13% | 60% |
|  | G45 | 3.26 | 0.82 | 25% | 0.24 | 0.031 | 0.05 | 0.32 | 1.1 | 39% | 35% |
|  | G54 | 3.26 | 1.2 | 36% | 0.51 | 0.038 | 0.10 | 0.64 | 1.8 | 55% | 56% |
|  | G63 | 3.26 | 1.8 | 55% | 0.034 | 0.011 | 0.77 | 0.82 | 2.6 | 45% | 81% |
|  | G14 +TCE | 3.26 | 1.0 | 32% | 0.62 | 0.095 | 0.20 | 0.91 | 2.0 | 87% | 60% |
| **AVERAGE originals** |  | **2.90** | **1.4** | **44%** | **0.30** | **0.06** | **0.22** | **0.57** | **2.0** | **40%** | **60%** |
| **Standard Dev (n-9)** |  | **0.72** | **0.39** | **11%** | **0.22** | **0.04** | **0.26** | **0.29** | **0.51** | **22%** | **14%** |
| TRANSFERS AMENDED REGULARLY | GT15 | 17.9 | 5.6 | 31% | 2.4 | 0.69 | 1.1 | 4.2 | 9.8 | 75% | 54% |
|  | GT3 | 15.8 | 5.7 | 36% | 4.1 | 0.11 | 1.4 | 5.6 | 11 | 98% | 71% |
|  | GT5 | 23.6 | 7.3 | 31% | 1.3 | 0.51 | 1.1 | 2.9 | 10 | 40% | 43% |
|  | GT20 | 25.7 | 8.4 | 33% | 2.7 | 1.0 | 1.1 | 4.8 | 13 | 58% | 52% |
|  | GT33 | 19.6 | 6.1 | 31% | 1.5 | 0.23 | 1.1 | 2.9 | 9.0 | 48% | 46% |
|  | GT4 | 22.4 | 6.0 | 27% | 0.48 | 0.16 | 0.63 | 1.3 | 7.2 | 21% | 32% |
| **AVERAGE transfers** |  | **20.8** | **6.5** | **31%** | **2.1** | **0.45** | **1.1** | **3.6** | **10** | **57%** | **50%** |
| **Standard Dev (n-6)** |  | **3.72** | **1.1** | **3%** | **1.3** | **0.35** | **0.24** | **1.6** | **2.0** | **27%** | **13%** |

Using these estimated response factors applied to the areas determined by LC/MS analysis of well-mixed slurry samples, we could calculate total moles recovered per bottle by multiplying concentrations by the total slurry volume. We then compared the CLD and metabolite moles recovered in the bottles after 8 years to the initial amount of CLD added (Table 2; calculations in Table S10). The three different groups of microcosms, poisoned controls, active original microcosms, and active microcosm transfers, did indeed show differences in CLD recovery. In the group of seven poisoned controls, two microcosms (G1 and G3) produced a lot of methane and thus were biologically active, despite having been poisoned. These bottles also exhibited extensive metabolite production, unlike remaining controls (Figure 1 and Table 2). Therefore, for the mole balance analysis, we included those two microcosms into the group of active original microcosms. We were able to recover 63±6% of added CLD in the poisoned controls after 8 years, 44±11% in the original microcosms that only receive electron donor in the first 2 years, and only 31±3% of added CLD in the transfers amended regularly with donor and CLD (Table 2). The loss of ~37% in the poisoned control group likely results from sorption to glass and stoppers, poor extraction from soils during sample preparation, losses from volatilization, and some minimal losses (<1%) from sample removal. Losses in microbially-active bottles are greater and can be explained by the contribution of biological transformation processes.

We estimated the total moles recovered as metabolites in the various treatments. Metabolites were not detected in the un-amended slurry microcosms, nor in the medium controls. The inactive poisoned controls had only trace amounts of MHCLD (0.001-0.004 μmoles) and no other metabolites. The active bottles had significantly higher concentrations of metabolites, especially in the transfers that received more CLD and donor. The estimated sum of moles of metabolites ranged from 13% to 98% of the CLD remaining after 8 years in active microcosms. When the sum of all measured metabolites was included in the mole balance, overall recoveries after 8 years were more similar regardless of treatment, 63±6%, 60±14%, and 50±13% for controls, originals and transfers, respectively. Given the length of the study and approximations in calibration factors, these results were very reassuring and provided confidence in the measurements.

### Microbial Community Analysis qPCR and Sequencing Analysis

While DNA samples were collected and analyzed at various times throughout the 8 years, preliminary analyses have not revealed any clear trends to date. A snap shot of the microbial community and abundance after 77 months is shown in Figure 4 (raw data are provided in Tables S11 and S12). The microbial community of the 6 transfers reveals an abundance of fermentative and syntrophic anaerobes, consistent with the electron donor mix (acetone and ethanol) provided. Microcosms GT33 and GT4 produced little to no methane, and contain few or no methanogens. GT4 is the most extensively transferred microcosm and no longer contains soil. It also has the lowest bacterial cell numbers inferred from qPCR of 16S rRNA copies per ml. No particular trends are discernable at this time. Perhaps, as concluded by Chaussonnerie et al., [32], the microbial transformation is cometabolic, and really only dependent on sufficient availability of reduced vitamin B12. Further studies are clearly warranted.

**Figure 4:**
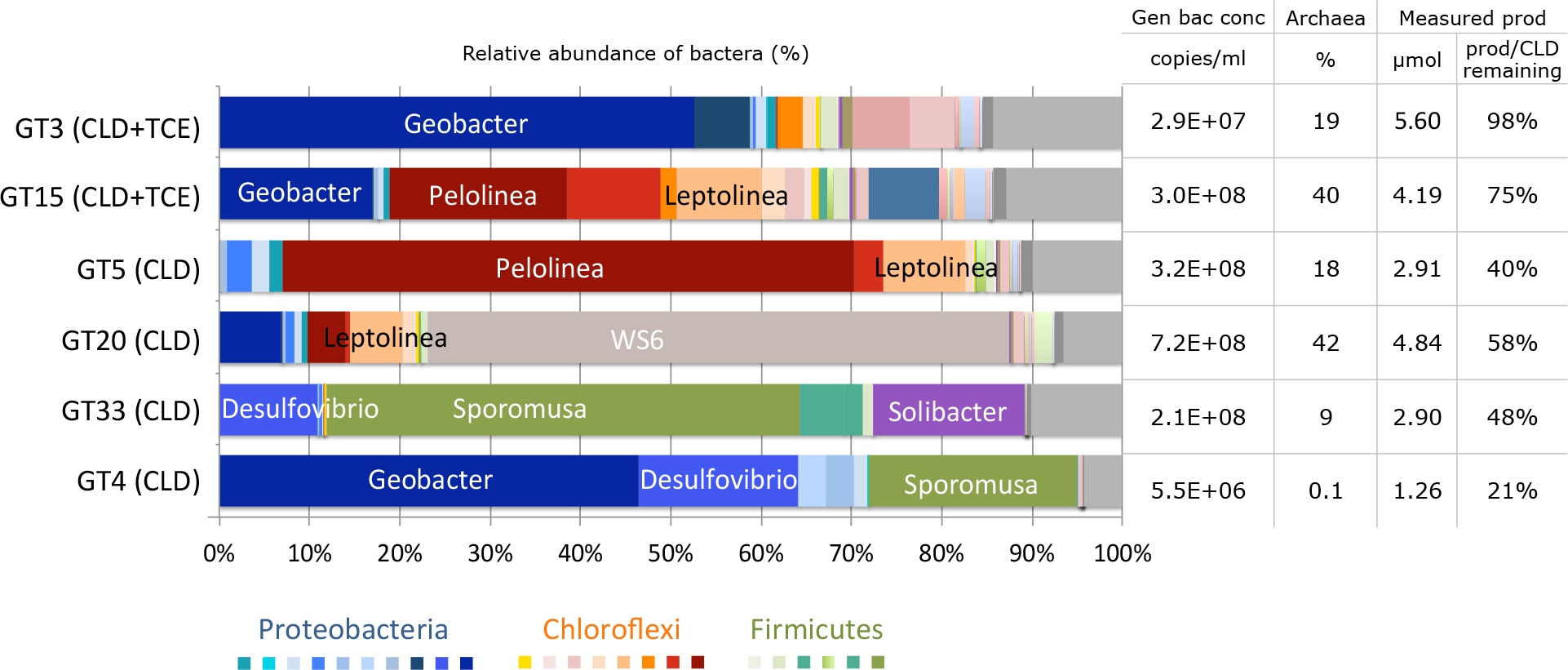
Microbial community in microcosm transfers. DNA was sampled 77 months after microcosm setup. The bar chart to the left shows relative abundance of bacteria obtained through small subunit (SSU) rRNA gene fragment sequencing. Table to the right shows concentration of general bacteria measured by qPCR (copies/ml), relative abundance of archaea (%) from sequencing, and measured products (estimated) (total, μmol and product/CLD remaining, %) for each bottle. Raw data of sequencing results and qPCR measurements can be found in Table S11 and S12.

### Analysis of Field Samples from 2018

We wondered if metabolites identified in the microcosms could also be detected in field samples, therefore we collected fresh soil sampless from the same locations in Guadeloupe that were previously sampled for the microcosm study and 9 samples were analyzed. (Table 3; raw data in Table S13). Anticipating quite low concentrations in the field samples, we decided to sample larger 5 and 20 ml volumes and perform a liquid-liquid extraction with a concentration step (method 5 and 6, SMD 2) in addition to our already established 1 ml slurry sample preparation method (method 4, SMD 2). Analysis of 6 soil samples from banana plantations in Guadeloupe revealed CLD concentrations in the range 120 to 1000 ng/g soil, or 0.12 to 1.0 mg/kg. These values are pretty typical of Guadeloupe soils: a recent survey [42] reported CLD concentrations in mg/kg in soil of 0.03 (minimum), 2.00 (median), 3.39 (mean), and 24.2 (maximum). MHCLD was detected in all soil samples and ranged from 1 to 8% of CLD based on area from LC/MS analysis. Most soil samples also showed DHCLD, but area counts were 10 to 100 times lower than those of MHCLD. Pentachloroindene (B5a) concentrations were estimated based on response factor for B1, and ranged from 0.5 to 24 ng/g of solid. B4a metabolites were also detected in the soil samples, but area counts were generally lower (up to 30 times) than those of B5a. Two of the 3 activated carbon sludge samples showed quite high CLD concentrations, between 6,000 and 8,500 ng/g. MHCLD concentrations were also significant, between 4 and 10% of CLD based on area counts. No PCIN was detected in the activated carbon sludge samples. We did not detect CPCIN metabolites in any of the field samples. Due to their hydrophilicity, the CPCIN metabolites would likely not be captured using the liquid-liquid extraction/concentration method (methods 5 and 6) used, and we did not have an alternative method for concentration of these compounds. We could also hypothesise that due to their hydrophilic nature and low sorption, these compounds may have been washed out from the soils by rain. Regardless, these field data confirm that anaerobic ring opening and dechlorinating processes do occur *in situ* in Guadeloupe soils. A more extensive analysis is thus warranted to determine locations for highest intrinsic activity on the islands and if rates could be accelerated by inducing anaerobic conditions, such as through organic amendment. The mechanism of CLD biotransformation also needs further investigation.

**Table 3:** Chlordecone and dechlorinated metabolites detected in field samples taken in 2018 from Guadeloupe. No carboxylated polychloroindenes were detected in these samples. Numbers are maximum area from the LC/MS analysis of two different slurry sample sizes (5 and 20 ml). Raw data in Table S13. CLD, MHCLD and DHCLD are quantified/reported as CLD hydrate, MHCLD hydrate and DHCLD hydrate.

|  | Chlordecone |  | Hydrochlordecones |  | Polychloroindenes |  |  |
| --- | --- | --- | --- | --- | --- | --- | --- |
|  | CLD |  | MHCLD (A9a) | DHCLD (A8a) | Pentachloroindene (B5a) |  | Tetrachloroindene (B4a) |
|  | area/g solids | ng/g solids | area/g solids | area/g solids | area/g solids | ng/g solids* | area/g solids |
| AgSoil1 | 1.4E+07 | 120 | 8.7E+05 | 2.1E+04 | 6.3E+03 | 0.48 | 4.1E+03 |
| AgSoil2 | 2.7E+07 | 130 | 1.9E+06 | 1.6E+04 | 3.0E+04 | 2.3 | 5.9E+03 |
| AgSoil3 | 1.9E+08 | 1000 | 5.3E+06 | 8.0E+04 | 3.1E+04 | 2.3 | 9.9E+02 |
| AgSoil4 | 6.2E+07 | 270 | 5.1E+06 | 8.8E+04 | 3.1E+05 | 24 | 4.2E+04 |
| AgSoil5 | 5.7E+07 | 130 | 4.2E+06 | 2.2E+04 | 1.1E+05 | 8.8 | 2.6E+04 |
| AgSoil6 | 3.5E+07 | 430 | 5.0E+05 | 0.0E+00 | 1.5E+04 | 1.1 | 1.0E+04 |
| AC-1 | 1.5E+07 | 24 | 1.5E+06 | 7.0E+05 | ND | ND | ND |
| AC-2 | 3.6E+09 | 6000 | 2.8E+08 | 1.8E+07 | ND | ND | ND |
| AC-3 | 5.1E+09 | 8400 | 1.9E+08 | 5.8E+06 | ND | ND | ND |

\*estimated values based on analysis of B1 standards run in 2019
ND: Not detected

## CONCLUSIONS

We have provided convincing LC/MS evidence for extensive dechlorination of CLD by indigenous microorganisms in chlordecone polluted soils. At least 19 different metabolites were detected as CLD concentrations progressively decreased over long-term microcosm incubations. Metabolites included hydrochlordecones, and the open-cage polychlorinated indenes and polychlorinated carboxylated indenes. Evidence for up to 9 Cl removed from the parent chlordecone molecule was found. Carboxylated intermediates were found to sorb poorly to soil. They may be flushed away while the polychlorinated indenes stick strongly to the soil. Further experiments are warranted to determine how to increase dechlorination rates and to further study the fate of these new CLD metabolites. The good news is that less chlorinated open-cage structures are more likely to be biodegradable by a wider variety of microbes under both aerobic and anaerobic conditions; this is the first glimpse of hope that anaerobic bioremediation may be a viable approach for chlordecone.

## Abbreviations

CLD: Chlordecone
MHCLD: Monohydrochlordecone
DHCLD: Dihydrochlordecone
THCLD: Trihydrochlordecone
HCLD: Hydrochlordecone
PCIN: Polychloroindene
CPCIN: Carboxylated polychloroindene
TCE: Trichloroethene
VC: Vinyl Chloride
MC: Microcosm
GC: Gas Chromatograph
FID: Flame Ionization Detector
IC: Ion Chromatograph
LC: Liquid Chromatograph
MS: Mass Spectrometry
eeq: Electron Equivalents
SMD: Supplemental Method Details

## ASSOCIATED CONTENT

### Supporting Information

Suporting Method Details (SMD) for field sampling and for microcosm setup including recipe for artificial groundwater, analysis procedure and sample preparation methods for GC-FID, IC, pH and LC/MS analysis, and DNA extraction, amplicon sequencing, and qPCR analysis procedures. Figures showing microcosm transfers and their origins, history of microcosms (CLD added, donor added, methane produced), methane production in active CLD amended MCs during the first 1.5 years of monitoring, TCE and its dechlorination products in a CLD and TCE amended MC (G15) during the first 1.5 years of monitoring, LC/MS scan analysis of CLD (CLD hydrate, hemi-acetal and other observed CLD adducts), and chromatograms and spectra for each of the 19 observed metabolites. Tables showing Guadeloupe field sample details, treatment table for Guadeloupe microcosm study, details of feeding stock preparation, details of CLD and donor amendments, details of GC-FID and IC analysis and pH measurements, list of dechlorinated metabolites identified in microcosm samples, results of all LC/MS analysis, normalized LC/MS area counts in GT20, sorption calculations and plots, mass balance calculations, sequencing analysis and qPCR analysis, and concentration calculations for CLD and its degradation products in Guadeloupe field samples.

## ACKNOWLEDGEMENTS

We thank Melanie Duhamel (U of Toronto) and Emmanuel Duquesnoy (SIAEAG) for initiating the collaboration between Guadeloupe and Toronto in 2010. We also thank Marion Chevallier, Pierre-Loïc Saaidi and Denis Le Paslier (Université d'Évry Val-d'Essonne & Genoscope, France) for providing us with a sample of their purified metabolite B1, and for very helpful discussions regarding metabolite identification. Post-doctoral and PhD stipends were provided by Region Guadeloupe-FEDER 2007-2013. Funding was provided by the Government of Canada through Genome Canada and the Ontario Genomics Institute (2009-OGI-ABC-1405 and OGI-102), from the Natural Science and Engineering Research Council (NSERC) of Canada, and from the Ontario-China Research and Innovation Fund (OCRIF).

