## Supplemental information text file for "Evidence for extensive anaerobic dechlorination and transformation of the pesticide chlordecone (C_10_Cl_10_O) by indigenous microbes in microcosms from Guadeloupe soil"

### TABLE OF CONTENTS

### SUPPLEMENTAL METHOD DETAILS (SMD)


1. Field sampling and microcosm setup
    A) Collection of field samples
    B) Recipe for artificial groundwater used in microcosm setup
2. Microcosm sampling and analysis
    A) GC-FID sample preparation and analysis
    B) IC sample preparation and analysis
    C) pH measurements
    D) LC/MS sample preparation and analysis
    E) DNA extraction, amplicon sequencing and Quantitative Polymerase Chain Reaction (qPCR) analysis, including modifications to the DNeasy PowerSoil Kit manufacturer's protocol


### LIST OF SUPPORTING FIGURES


**Figure S1**: Microcosm transfers and their origins.

**Figure S2:** History of microcosms. Cumulative chlordecone added (a), cumulative donor added (b), and cumulative methane produced (c) in active CLD amended (GT20, GT5, GT33 and GT3) and active CLD+TCE amended bottles (GT15, GT3) over the course of the study

**Figure S3:** Methane production in active CLD amended microcosm during the first 1.5 years of monitoring.

**Figure S4:** TCE and its dechlorinated metabolites in a TCE and CLD amended microcosm during the first 2 years of monitoring (microcosm G15)

**Figure S5:** LC-MS scan analysis of chlordecone in standard run on June 29$^{th}$ 2018 (instrument method and standard preparation method is described in main text and in Supplemental Method Details 2) (a) m/z 502.6874 EIC (extracted ion chromatogram), (b) m/z 548.6929 EIC, (c) m/z 562.7078 EIC, (d) m/z 516.7032 EIC, (e) full scan mass spectra at 9.9 min,

**Figure S6 A-J:** Chromatograms and spectra for each of the 19 observed metabolites. (A): Monohydrochlordecone A9a, (B): Dihydrochlordecone A8a, (C): Trihydrochlordecone A7a, (D): Pentachloroindene B5a, (E): Tetrachloroindene B4a, (F): Trichloroindene B3a, (G): Carboxylated tetrachloroindenes C4a-b, (H): Carboxylated trichloroindenes C3a-e, (I): Carboxylated dichloroindenes C2a-d, (J): Carboxylated chloroindenes C1a-b.

### LIST OF SUPPORTING TABLES

*All Supporting information tables are provided in a spreadsheet format in the accompanying Excel file*

**Table S1 A and B:** Guadeloupe field sample details. (A): Field samples used for microcosm setup, (B): Field samples analyzed by LC/MS

**Table S2 A-C:** Treatment table for Guadeloupe microcosm study. (A): Microcosms, (B): Transfers, (C): Medium controls

**Table S3:** Details of feeding stock preparation for the microcosm study

**Table S4:** Details of chlordecone and donor amendments, DNA sampling times and CLD analysis dates in microcosms and transfers over time

**Table S5:** Details of GC-FID analysis (methane, ethene, VC, trans-DCE, cis-DCE and TCE), IC analysis (acetate, chloride, nitrite, sulfate, nitrate, phosphate) and pH measurements in microcosms and transfers over time

**Table S6:** Chlordecone and dechlorinated metabolites identified in microcosm samples

**Table S7:** Results from LC/MS analysis of samples analyzed over the course of the study

**Table S8:** Normalized LC/MS area counts of metabolites in transfer GT20 over the course of the study

**Table S9:** Calculating sorption of CLD and its metabolites to soil in microcosms and transfers

**Table S10:** Mass balance calculations for CLD and its dechlorinated metabolites in some microcosms and transfers

**Table S11:** Data from Illumina MiSeq sequencing

**Table S12:** Data from qPCR analysis

**Table S13:** Concentration calculations for CLD and its dechlorinated metabolites in Guadeloupe field samples

### SUPPLEMENTAL METHOD DETAILS (SMD)

#### 1. Field Sampling and Microcosm Setup

**A) Collection of Field Samples**

All agricultural soil samples were collected from 0-20 cm depth. In 2010, soil, water and sludge samples were collected separately in polypropylene sampling bottles, while in 2018, a mix of soil and water was placed into 1-liter glass sampling jars that were filled to the top and sealed. All samples were shipped from Guadeloupe to University of Toronto for analysis where they were stored at 4°C until use.

**B) Recipe for Artificial Groundwater Used in Microcosm Setup**

From P.P.J. Middeldorp et al. “Stimulation of reductive dechlorination for in situ bioremediation of a soil contaminated with chlorinated ethenes.” Water Science Technology Vol 37. No. 8, pp. 105-110. 1998.

For 1 liter of water, the following compounds were added:

| Chemical | Concentration | Molecular weight | Amount required |
|---|---|---|---|
| | (mM) | (g/mol) | (mg) |
| $NH_4Cl$ | 1 | 53.49 | 53.5 |
| $MgCl_2$ (6 $H_2O$) | 0.05 | 203.31 | 10.2 |
| $MnCl_2$ (4 $H_2O$) | 0.02 | 197.9 | 4.0 |
| NaCl | 0.12 | 58.44 | 7.0 |
| $CaCl_2$ | 6 | 147.02 | 882.1 |
| $Na_2HPO_4$ | 0.45 | 141.96 | 63.9 |
| $KH_2PO_4$ | 0.15 | 136.09 | 20.4 |
| $Na_2SO_4$ | 1 | 142 | 142.0 |
| $NaHCO_3$ | 1 | 86.01 | 86.0 |

The solution was made with distilled deionized water. It was autoclaved and then purged with $N_2/CO_2$ (80% /20%) for 1 hour. After, the artificial groundwater was moved into the glovebox. In the glovebox 10 mL of 100x sterile vitamins solution was added (recipe in Edwards, E. A. and D. Grbic-Galic, 1994 "Anaerobic Degradation of Toluene and o-Xylene by a Methanogenic Consortium." Applied and Environmental Microbiology 60(1): 313-322). The pH was measured and adjusted to pH 7.

### 2. Microcosm Sampling and Analysis

#### A) GC-FID Sample Preparation and Analysis

Microcosms were sampled using glass syringes and a 1 ml sample was added to a 10 ml headspace autosampler vials (Agilent) containing 5 ml of acidified deionized water (pH 2). The vials were then sealed with Teflon-coated septa and aluminum crimp caps (Chromatographic Specialties) for automated headspace injection onto the CG. Methane, ethene, ethane, and chlorinated ethene measurements were carried out as headspace analysis using an Agilent 7890A gas chromatograph (GC) equipped with a flame ionization detector (FID), a G1888 headspace autosampler and a J&W GS-Q column (30 m x 0.53 mm) (Agilent, Santa Clara, CA, USA). Helium was the carrier gas (11 ml/min) and the oven program was as follows: Hold at 35°C for 1.5 min, increase to 100°C at the rate of 15°C/min, then increase to 185°C at the rate of 5°C/min, hold 10 min, then increase to 200 °C at the rate of 20°C/min, hold 10 min (total runtime 43.6 min). The GC had a packed inlet and a 3 ml sample loop. Headspace operating temperatures for oven, loop and transfer line were 70, 80 and 90°C respectively, while vial equilibration time, pressurization time, loop fill time, loop equilibration time and injection time were 40, 0, 0.2, 0 and 3 min respectively. During equilibration in oven, the samples were shaken at low speed. Data from the GC was integrated using ChemStation (Agilent). Calibration standards were prepared in the concentration range 0.2 to 2 mg/l (liq) for methane and ethene, and 1 to 20 mg/l for chlorinated ethenes.

#### B) IC Sample Preparation and Analysis

To measure chloride, nitrite, nitrate, sulfate, phosphate and acetate concentrations, samples (1 ml) were filtered through 0.2 µm nylon filters (Fisher Scientific). Analysis were performed using a Dionex ICS 2100 ion chromatograph with a Dionex IonPac AS18 analytical column (4x250 mm) and an ASRS 500 suppressor (Thermo Fisher Scientific). The samples were run isocratically at 23 mM KOH, 57 mA current and with a flow rate of 1 ml/min. Data from the IC was integrated using Chromelion (Thermo Fisher Scientific). Standards were prepared in the range 0.005 to 0.5 mM.

#### C) PH Measurements

One ml liquid samples were taken for "in-syringe" pH analysis using an Oakton pH spear (Oakton Instruments, Vernon Hills, USA). If needed, pH was adjusted with Na-bicarbonate or HCl to pH 7.

### D) LC-MS Sample Preparation and Analysis

*Summary of sample preparation methods and LC-MS methods on the different analysis dates:*

| Analysis date | Sample prep method | Scan Range | Instrument | Eluent | Flow rate | Column Temp | Gradient |
|---|---|---|---|---|---|---|---|
| 02-Oct-13 | Method 1 | 140-520 | Thermo Exactive LCMS | Water (A)/Acetonitrile (B) gradient | 0.2 ml/min | Ambient | 50%B start, increase to 100%B over 25min, decrease to 50% B over 5min |
| 27-Aug-14 | Method 1 | 130-550 | Thermo Exactive LCMS | Water (A)/Acetonitrile (B) gradient | 0.2 ml/min | Ambient | 50%B start, increase to 100%B over 25min, decrease to 50% B over 5min |
| 24-Nov-16 | Method 2 * | 120-1800 | Thermo Q-Exactive LCMS | Water (A)/Methanol (B) gradient | 0.2 ml/min | 40ºC | 50%B start, increase to 100%B over 15min, hold for 9min, decrease to 50%B over 1min, hold for 5min |
| 02-Mar-17 | Method 3 | 150-750 | Thermo Q-Exactive LCMS | Water (A)/Methanol (B) gradient | 0.3 ml/min | 40ºC | 50%B start, increase to 100%B over 8min, hold for 4min, decrease to 50%B over 1min, hold for 5min |
| 15-Feb-18 | Method 3 | 150-750 | Thermo Q-Exactive LCMS | Water (A)/Methanol (B) gradient | 0.3 ml/min | 40ºC | 50%B start, increase to 100%B over 8min, hold for 4min, decrease to 50%B over 1min, hold for 5min |
| 06-Apr-18 | Method 4 | 150-750 | Thermo Q-Exactive LCMS | Water (A)/Methanol (B) gradient | 0.3 ml/min | 40ºC | 50%B start, increase to 100%B over 8min, hold for 4min, decrease to 50%B over 1min, hold for 5min |
| 09-Apr-18 | Method 4 | 150-750 | Thermo Q-Exactive LCMS | Water (A)/methanol (B) gradient w/5 mM ammonium acetate | 0.3 ml/min | 40ºC | 50%B start, increase to 100%B over 8min, hold for 4mn, decrease to 50%B over 1min, hold for 5min |
| 29-Jun-18 | Method 3, 4 and 5 | 150-750 | Thermo Q-Exactive LCMS | Water (A)/methanol (B) gradient w/5 mM ammonium acetate | 0.3 ml/min | 40ºC | 50%B start, increase to 100%B over 8min, hold for 4min, decrease to 50%B over 1min, hold for 5min |
| 25-Sep-18 | Method 3, 4 and 6 | 150-750 | Thermo Q-Exactive LCMS | Water (A)/methanol (B) gradient w/5 mM ammonium acetate | 0.3 ml/min | 40ºC | 50%B start, increase to 100%B over 8min, hold for 4min, decrease to 50%B over 1min, hold for 5min |
| 07-Mar-19 | Method 4 | 150-750 | Thermo Q-Exactive LCMS | Water (A)/methanol (B) gradient w/5 mM ammonium acetate | 0.3 ml/min | 40ºC | 50%B start, increase to 100%B over 8min, hold for 4min, decrease to 50%B over 1min, hold 5min |

* For this run, samples were contained in plastic well plates with slit rubber mats. Compounds may have sorbed to the plastic, and standards had evaporated (slit mats do not seal well). For all other runs, samples were contained in glass vials with Teflon lined septa.

*Details of the different sample preparation methods*

**Sample preparation method 1 (water phase with small amount of soil, liq/liq extraction):**
1) Let soil settle in bottles overnight
2) Sample 2 mL liquid from microcosms
3) Extract with 15/85 % acetone/hexane, 2 cycles of extraction, each w/ 5mL of acetone/hexane
4) Filter the solvent phase through hydrophobic filter into glass vial
5) Evaporate filtrate to dryness and re-dissolve in 2mL of MeOH

**Sample preparation method 2 (water phase):**
1) Let soil settle in bottles overnight
2) Sample 1 mL liquid from microcosms
3) Filter sample into new glass vial through 0.2 μm PTFE syringe filter

**Sample preparation method 3 (water phase):**
1) Let soil settle in bottles overnight
2) Sample 0.75mL liquid from microcosms, avoiding getting any soil into the sample, and transfer to glass vial
3) Centrifuge @ 3000rpm for 5min
4) Transfer 0.5mL clear liquid from centrifuged samples into glass vials with 0.5mL MeOH, mix
5) Filter sample into new glass vial through 0.2 μm PTFE syringe filter

**Sample preparation method 4 (samples with soil):**
1) Shake bottle and sample 1mL slurry from microcosm
2) Add to glass vial containing 1mL MeOH, vortex, shake gently for 10min and let sit for about 30 min (or overnight)
3) Centrifuge @ 3000rpm for 5min
4) Filter sample into new glass vial through 0.2 μm PTFE syringe filter

**Sample preparation method 5 (extraction of field samples, 5 ml slurry):**
1) Shake bottle and sample 5mL slurry from sampling jar into a 15 ml glass centrifuge tube
2) Add 5 ml of a 15% acetone and 85% hexane mixture to the centrifuge tube
3) Shake gently for 10 min, let sample sit for 1 hour, shake for 1 min
4) Centrifuge @ 3000 rpm for 5min
5) Transfer the solvent phase into new glass vial
6) Filter sample into new glass vial through 0.2 μm PTFE syringe filter
7) Evaporate to dryness
8) Re-dissolve in 1 ml methanol

**Sample preparation method 6 (extraction of field samples, 20 ml slurry):**
1) Shake bottle and sample 5mL slurry from sampling jar into a 15 ml glass centrifuge tube (prepare 4 tubes with 5 ml sample for each field sample)
2) Add 5 ml of a 15% acetone and 85% hexane mixture to the centrifuge tube
3) Shake gently for 10min and let sample sit for 30 min
4) Centrifuge @ 3000 rpm for 5min
5) Transfer the solvent phase into new glass vial (combine solvent from the 4 tubes into one vial)
6) Repeat steps 2 to 5 one more time
7) Evaporate to dryness
8) Re-dissolve in 0.5 ml MeOH
9) Filter sample into new glass vial using 0.2 μm PTFE syringe filter

**E) DNA Extraction, Amplicon Sequencing and Quantitative Polymerase Chain Reaction (qPCR) Analysis**

Samples for DNA extraction were taken from the 6 transfers (GT5, GT20, GT33, GT4, GT15, GT3) about 7 years into the study for microbial community analysis. Slurry samples (1 mL) were taken, centrifuged at 10 000 rpm for 20 min, and cell pellets were frozen at -80$^{o}$C for future DNA extraction. DNA was extracted using the DNeasy PowerSoil Kit (Qiagen, Hilden, Germany) according to manufacturer's protocol with some modifications (see below). DNA concentrations were verified by a NanoDrop1000 Spectrophotometer (Thermo Fisher Scientific) and Qubit Fluorometric Quantitation (Thermo Fisher Scientific), and the extracts were stored at - 80$^{o}$C.

Microbial community composition of the samples was assessed by small subunit (SSU) rRNA gene fragment sequencing and Quantitative Polymerase Chain Reaction (qPCR) analysis. Modified versions of the universal SSU primer set 926f and 1392r (926f-modified: AAACTYAAAKGAATWGRCGG; 1392r-modified: ACGGGCGGTGWGTRC), targeting the V6−V8 variable region of the 16SrRNA gene from bacteria and archaea as well as the 18S rRNA gene in eukarya, was used. Primers were derived from previous publications (Ferris MJ, Muyzer G & Ward DM (1996) Denaturing gradient gel electrophoresis profiles of 16S rRNA-defined populations inhabiting a hot spring microbial mat community. *Appl Environ Microbiol* **62**: 340-346, and Engelbrektson, A., Kunin, V., Wrighton, K. C., Zvenigorodsky, N., Chen, F., Ochman, H. and Hugenholtz, P. 2010. Experimental factors affecting PCR-based estimates of microbial species richness and evenness. *The ISME journal* 4 (5): 642–647). The resulting amplicons were sequenced at Genome Quebec Innovation Centre using the Illumina MiSeq platform (Illumina Inc., San Diego, USA).

The paired end reads were processed in mothur v.1.39.5. The reads were assembled into contigs with default settings (i.e., allowed primer differences = 0, quality scores threshold = 20) and trimmed to a maximum length of 520bp with no ambiguous bases allowed. Unique sequences were aligned to the Silva v128 seed alignment provided at the mothur web site (https://www.mothur.org/), and only sequences that aligned to the V6−V8 variable region were kept. The sequences were further de-noised by pre-clustering the sequences allowing for up to 2 differences. Chimeras were then removed and the resulting sequences were classified taxonomically by comparing to the Silva v128.nr alignment also available at the mothur web site. Finally, the sequences were clustered into 0.03 OTUs using the cluster.split algorithm.

The abundance of total bacterial 16S rRNA genes in the microcosms was estimated by qPCR using a CFX96TM real-time PCR detection system, with a C1000 thermocycler (Bio-Rad Laboratories Inc., Hercules, CA, USA). DNA extracts were analyzed using the general bacterial 16S rRNA primers 055f (5'-ATGGCTGTCGTCAGCT-3') and 1392r (5'-ACGGGCGGTGTGTAC-3') (Ferris et al 1996). The 20 µl qPCR reactions were prepared in a PCR cabinet (ESCO Technologies, Gatboro, PA) and were made up by 10 µl of SsoFastTM EvaGreen® SuperMix (Bio-Rad Laboratories Inc., USA), 0.5 µl of each forward and reverse primers (10uM stock, making final concentration of 250 nM for both primers), 7 µl of UV treated UltraPure Distilled water (Invitrogen, Grand Island, NY, USA), and 2 µl of DNA extract diluted 10x. The qPCR cycle was as follows: 98°C for 2 min, 40 cycles of 98°C for 5 seconds and 55°C for 10 seconds, followed by an increase from 65°C to 95°C at 0.5°C increments over 10 seconds. Calibration curves for qPCR were prepared from serial dilutions of target-containing plasmids between $10^1$ and $10^8$ gene copies/ml. The general bacteria gene copy number detection limit was 1.1 E05 copies per ml.

***Modifications to the DNeasy PowerSoil Kit manufacturer's protocol***

Link to Dneasy PowerSoil kit Protocol:

https://www.qiagen.com/ca/resources/resourcedetail?id=91cf8513-a8ec-4f45-921e-8938c3a5490c&lang=en

Modifications to protocol:
Step 1: Cell pellet was added to the tube (instead of soil sample)
Step 9: All of the supernatant was transferred to the collection tube (not only 600 µl)
Step 12: All of the supernatant was transferred to the collection tube (not only 750 µl)
Step 19: Added 50 µl H2O instead of 100 µl C6

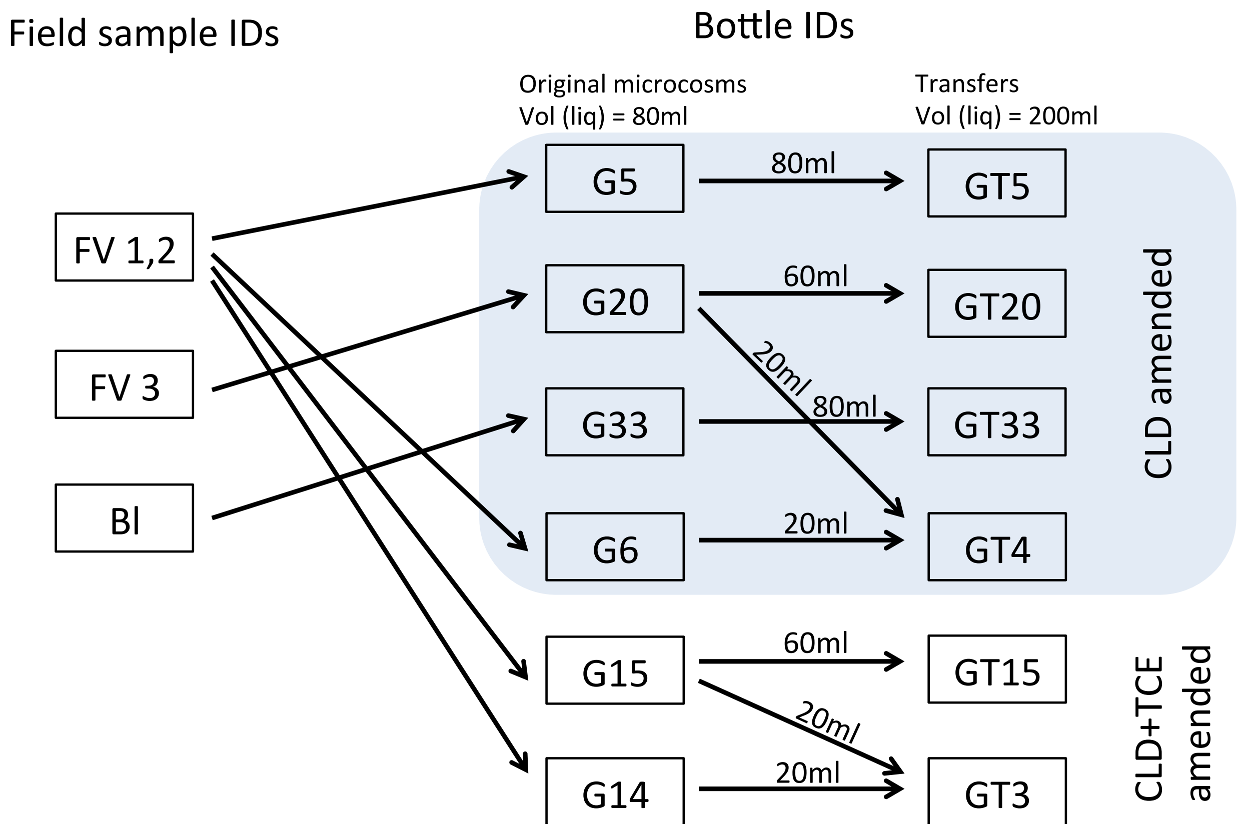


**Figure S1**: Microcosm transfers and their origins. Field samples FV 1, 2 and 3 were collected on a riverbank near a former banana field, and samples Bl inside an active agricultural banana production area in Guadeloupe

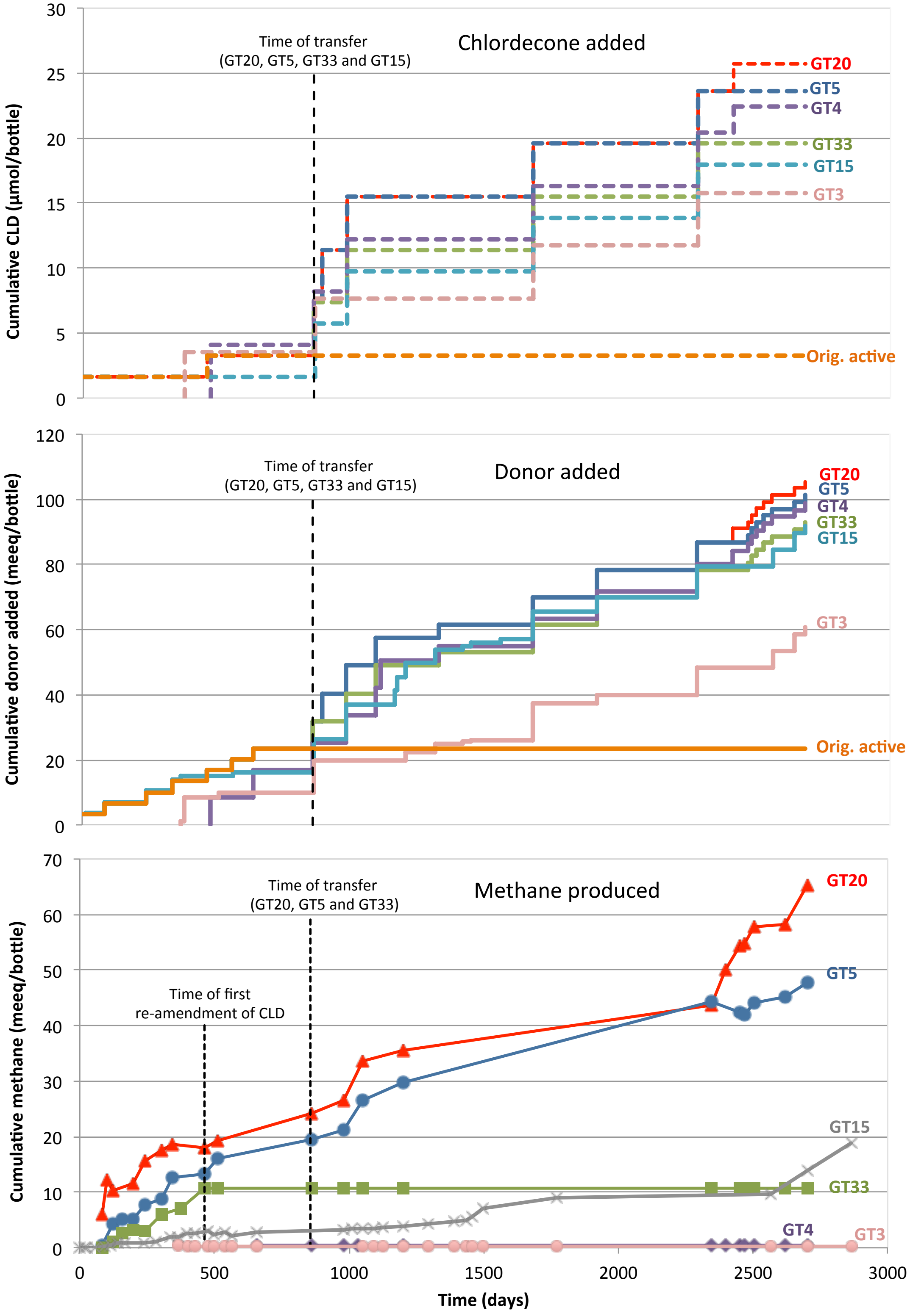


**Figure S2**: **History of microcosms.** Cumulative chlordecone added (a), cumulative donor added (b), and cumulative methane produced (c) in active CLD amended (GT20, GT5, GT33 and GT4) and active CLD+TCE amended bottles (GT15, GT3) over the course of the study. See Tables S4 and S5 for raw data and details of amendments and analyses.

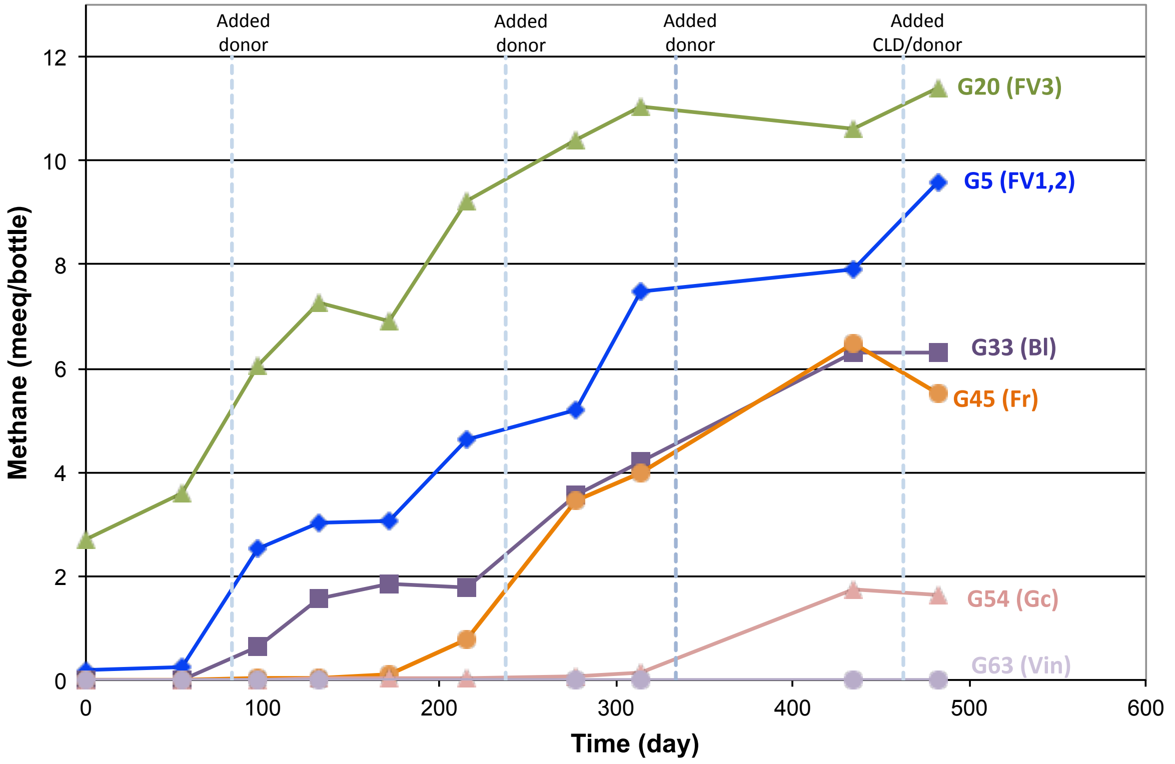


**Figure S3:** Methane production in active CLD amended microcosm during the first 1.5 years of monitoring. Data is shown for one of triplicates established for each soil type (soil type ID in brackets), but other replicates showed similar results.

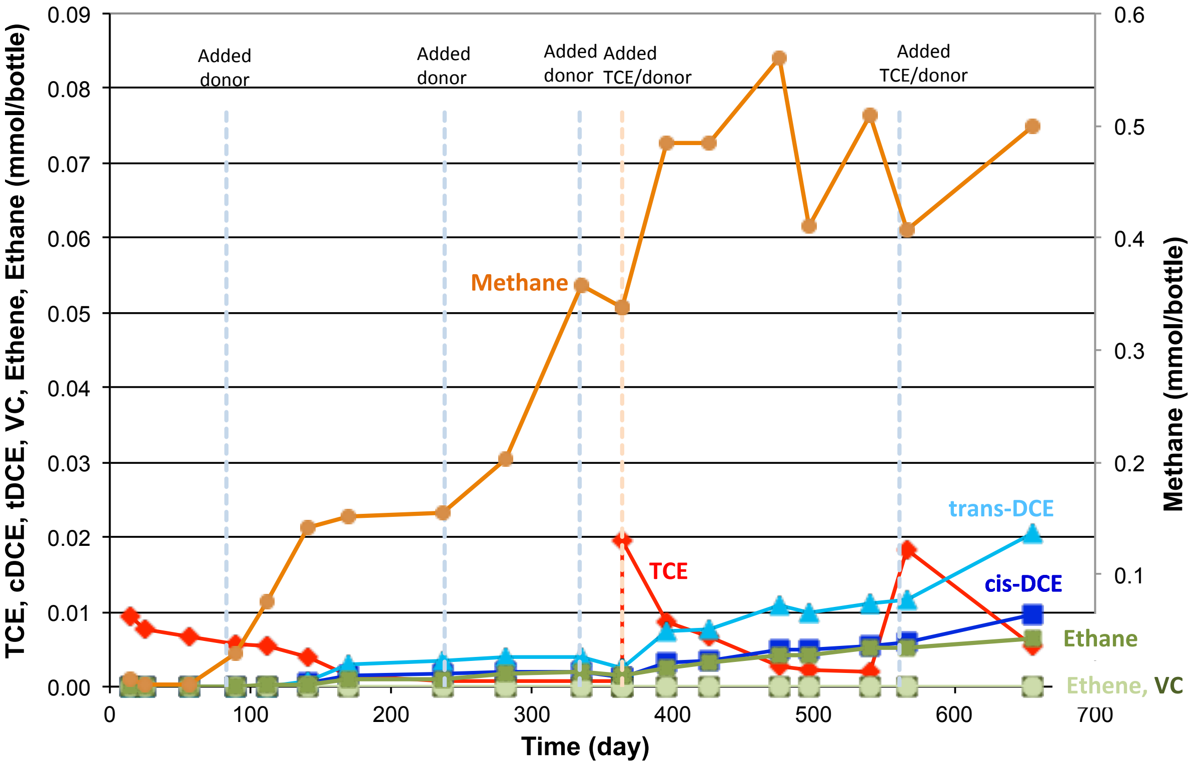


**Figure S4:** TCE and its dechlorinated metabolites in a CLD+TCE amended microcosm during the first 2 years of monitoring (microcosm G15). The two other triplicates (G13 and G14) behaved similarly. TCE continued degrading in G15 and after transferring in GT15, but GT3 which was a transfer from G14 and G15, never degraded TCE (Figure S2 and Table S5)

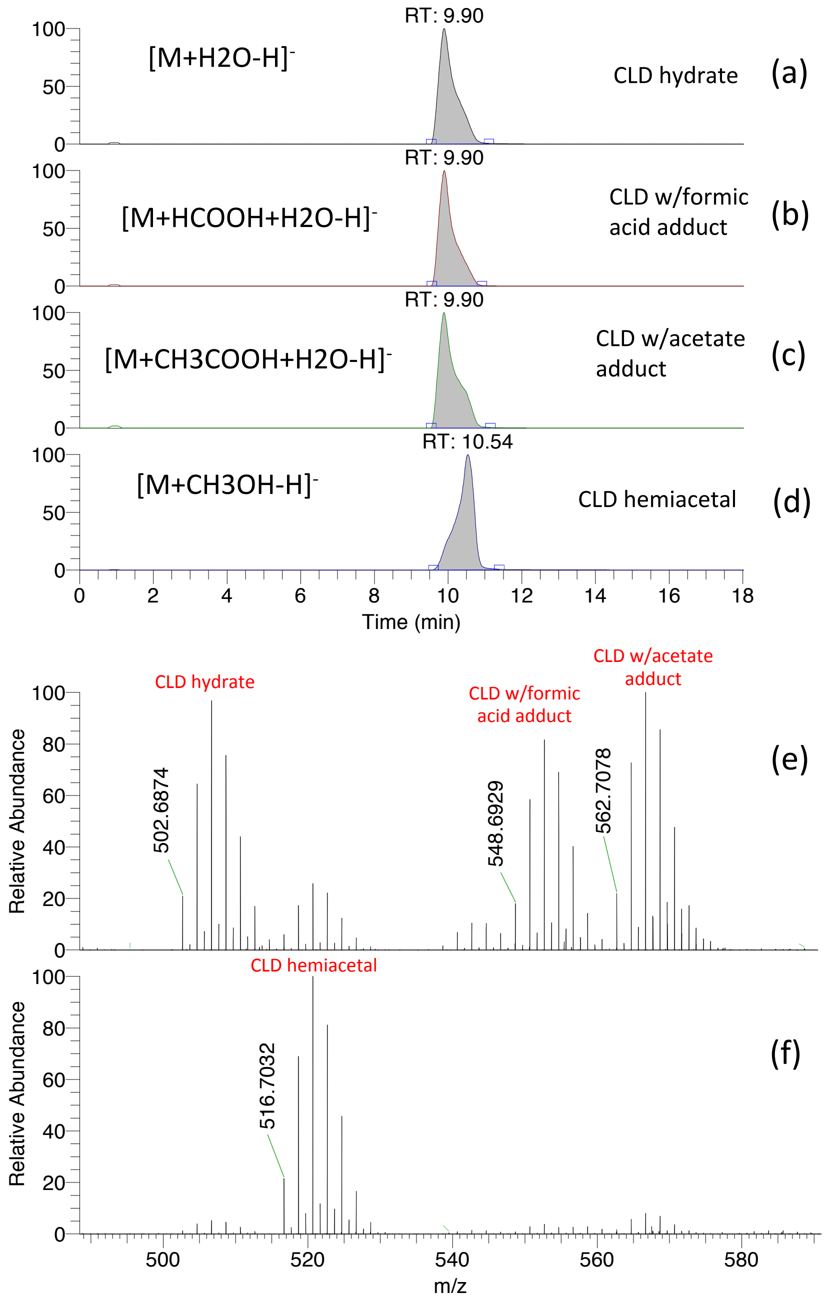


**Figure S5:** LC/MS scan analysis of chlordecone in standard run on June 29th 2018 (instrument method and standard preparation method is described in main text and in Supplemental Method Details 2) (a) m/z 502.6874 EIC (extracted ion chromatogram), (b) m/z 548.6929 EIC, (c) m/z 562.7078 EIC, (d) m/z 516.7032 EIC, (e) full scan mass spectra at 9.9 min, (f) full scan mass spectra at 10.54 min

Abbreviations:
NL: Normalized Level (average spectrum displayed normalized to the average base peak)
RT: Retention Time
AA: Peak area (automatically detected area)
AH: Peak height (automatically detected height)
m/z: An ion's mass-to-charge ratio

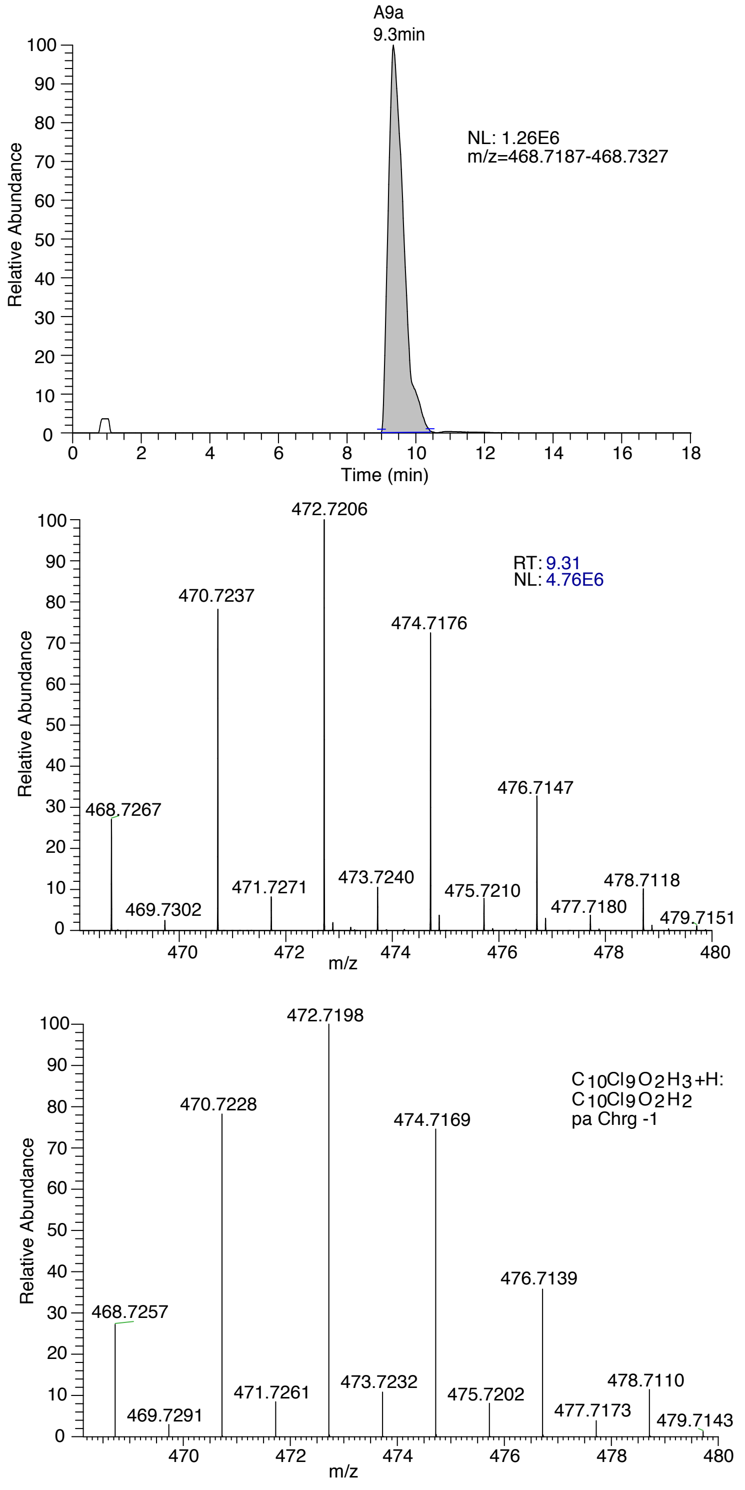


**Figure S6 A:** Extracted Ion Chromatogram, observed mass spectrum and theoretical mass spectrum for metabolite monohydrochlordecone (MHCLD) m/z 468.7267 (A9a, loss of 1 Cl), data from sample GT20 June 29th 2018. The mass tracked in this study is the hydrate form of MHCLD.

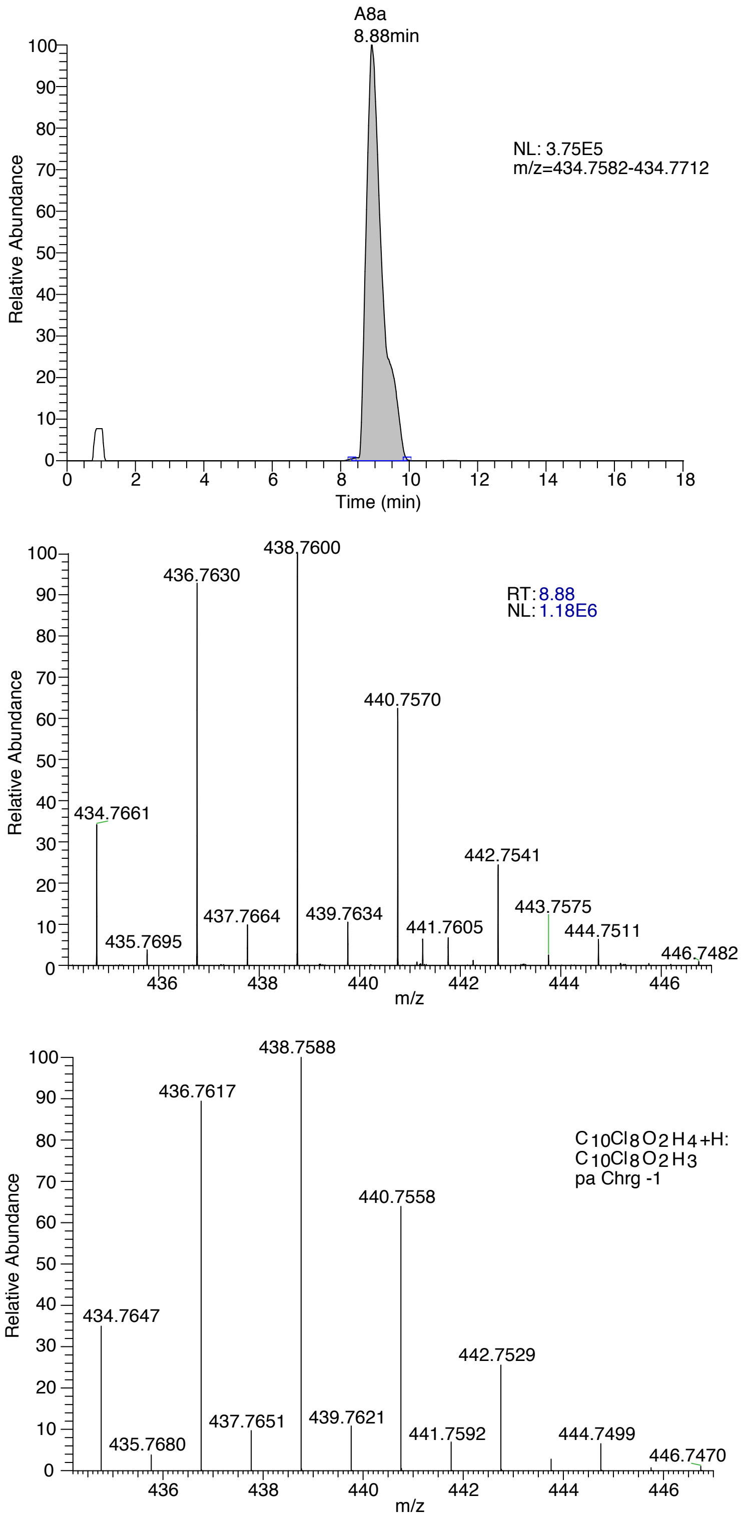


**Figure S6 B:** Extracted Ion Chromatogram, observed mass spectrum and theoretical mass spectrum for metabolite dihydrochlordecone (DHCLD) m/z 434.7661 (A8a, loss of 2 Cl), data from sample GT20 June 29th 2018. The mass tracked in this study is the hydrate form of DHCLD.

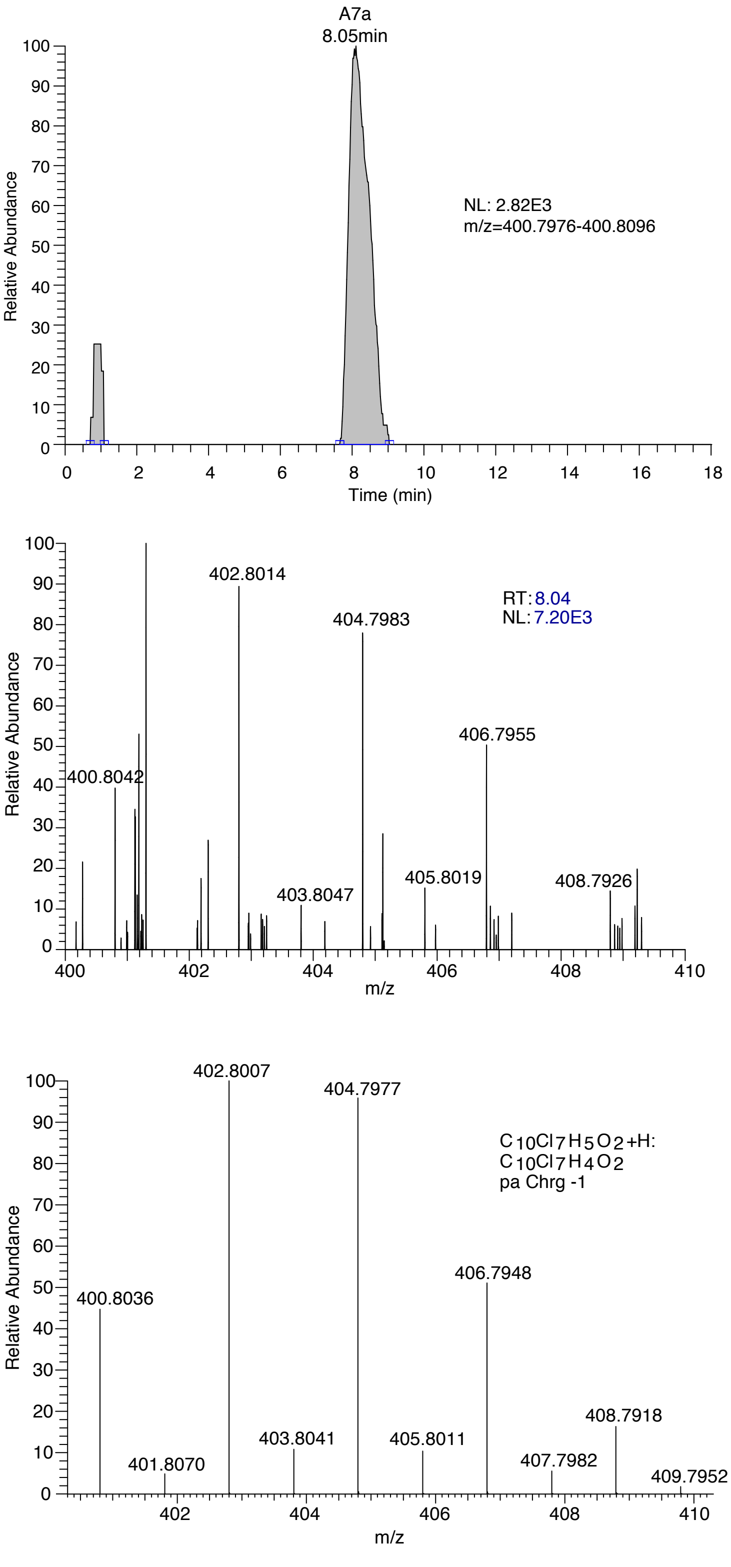


**Figure S6 C:** Extracted Ion Chromatogram, observed mass spectrum and theoretical mass spectrum for metabolite trihydrochlordecone (THCLD) m/z 400.8042 (A7a, loss of 3 Cl), data from sample GT3 March 7th 2019. The mass tracked in this study is the hydrate form of THCLD.

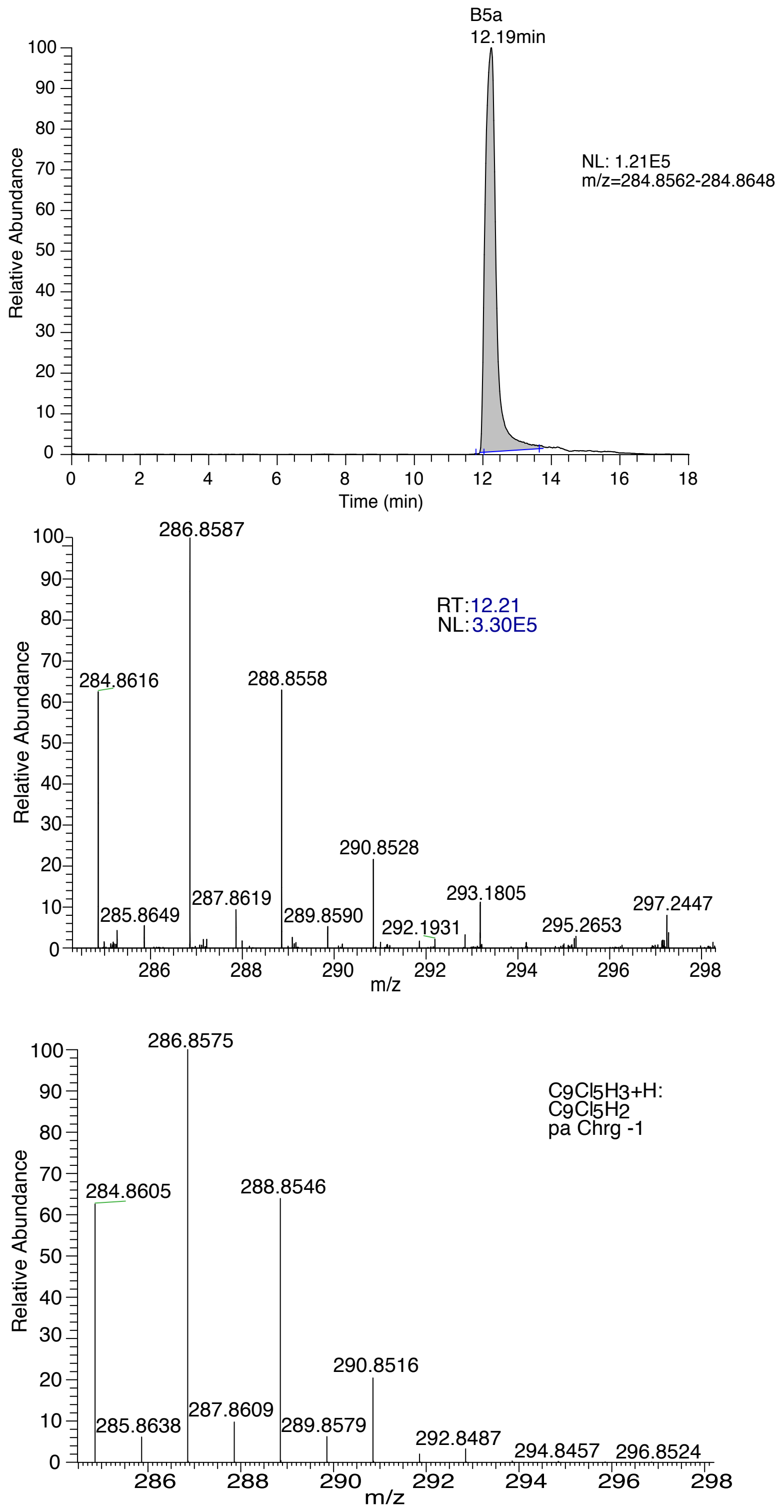


**Figure S6 D:** Extracted Ion Chromatogram, observed mass spectrum and theoretical mass spectrum for metabolite B5a m/z 284.8616 (pentachloroindene, loss of 5 Cl), data from sample GT20 June 29th 2018.

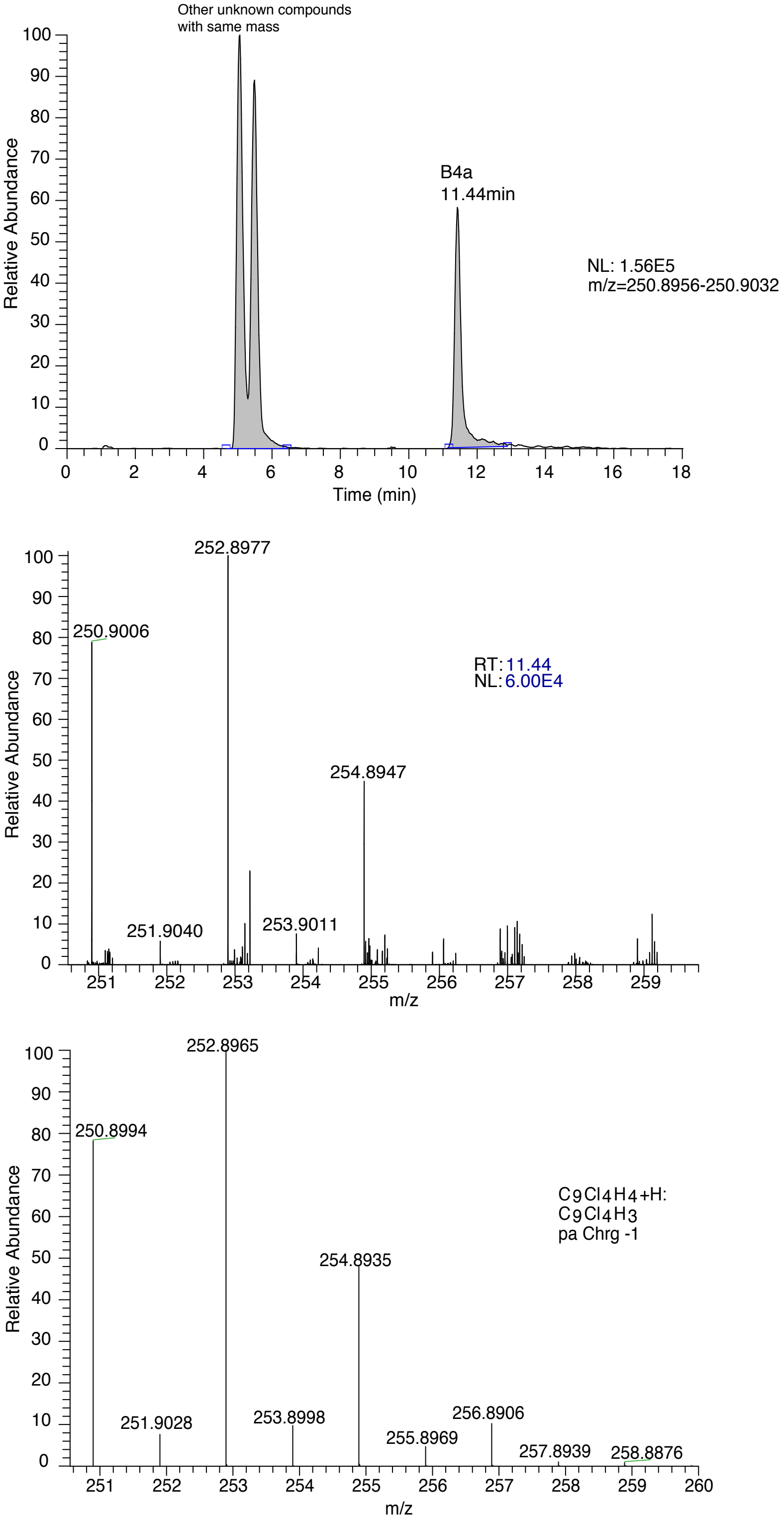


**Figure S6 E:** Extracted Ion Chromatogram, observed mass spectrum and theoretical mass spectrum for metabolite B4a m/z 250.9006 (tetrachloroindene, loss of 6 Cl), data from sample GT20 June 29th 2018.

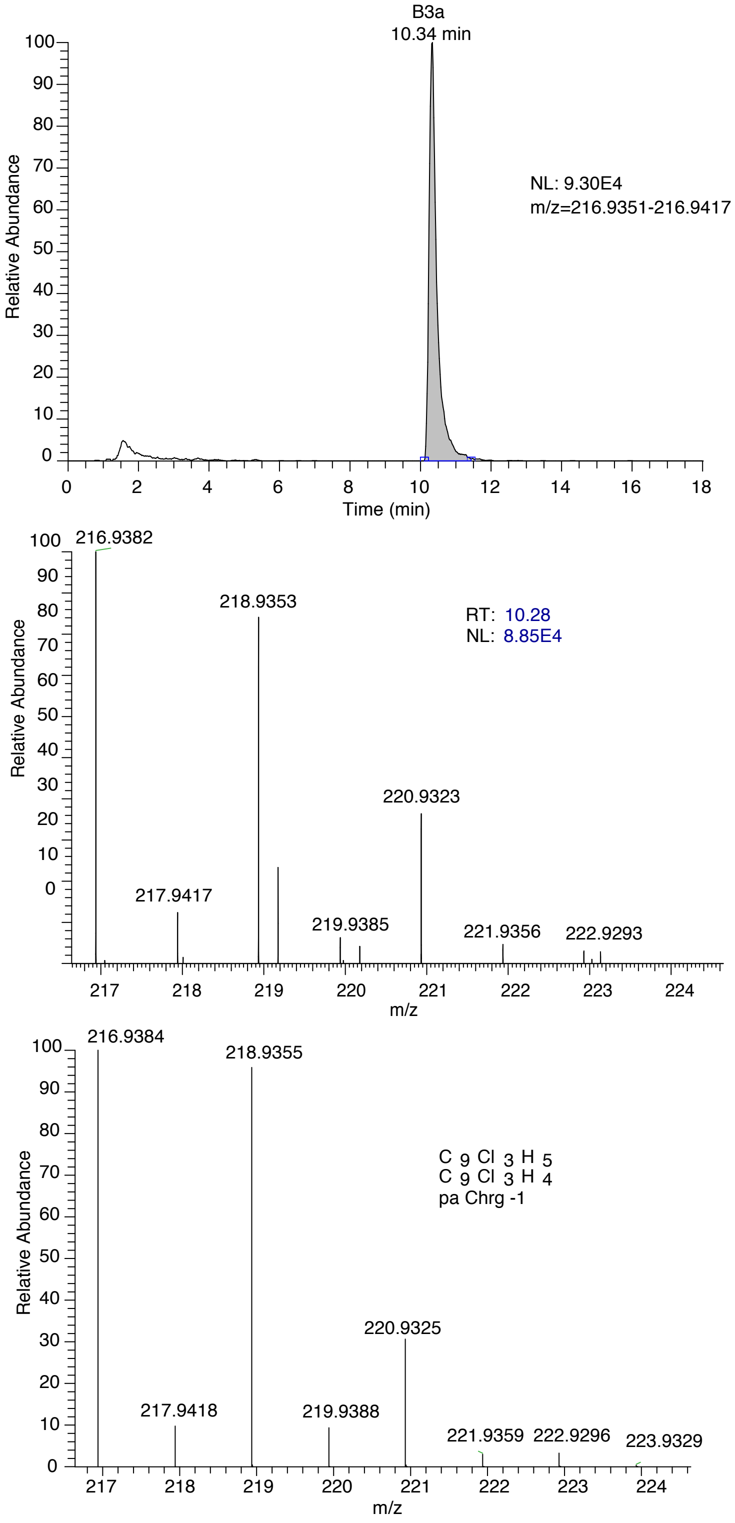


**Figure S6 F:** Extracted Ion Chromatogram, observed mass spectrum and theoretical mass spectrum for metabolite B3a m/z 216.9382 (trichloroindene, loss of 7 Cl), data from sample G19 March 7th 2019.

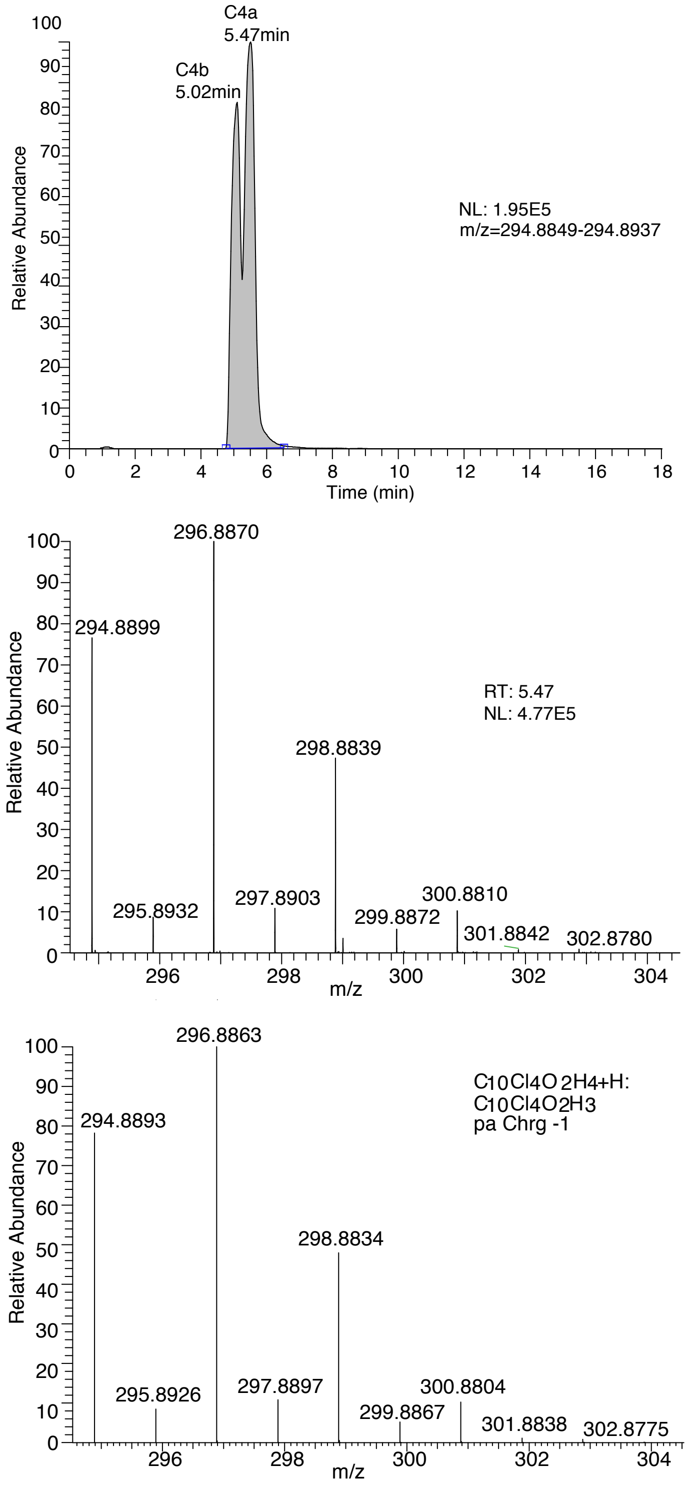


**Figure S6 G:** Extracted Ion Chromatogram, observed mass spectrum and theoretical mass spectrum for metabolites C4a-b m/z 294.8899 (carboxylated tetrachloroindene, loss of 6 Cl), data from sample GT20 June 29th 2018.

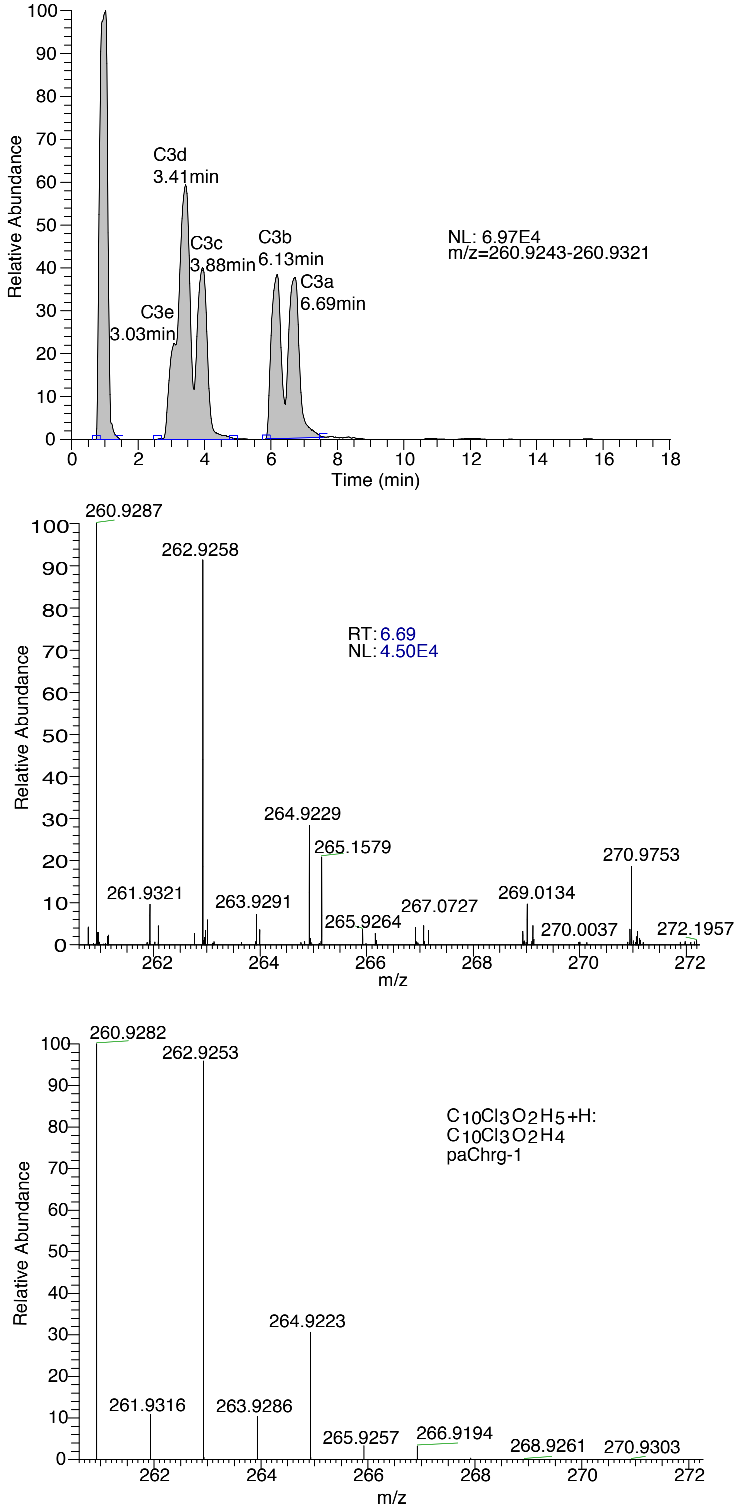


**Figure S6 H:** Extracted Ion Chromatogram, observed mass spectrum and theoretical mass spectrum for metabolites C3a-e m/z 260.9287(carboxylated trichloroindene, loss of 7 Cl), data from sample GT20 June 29th 2018.

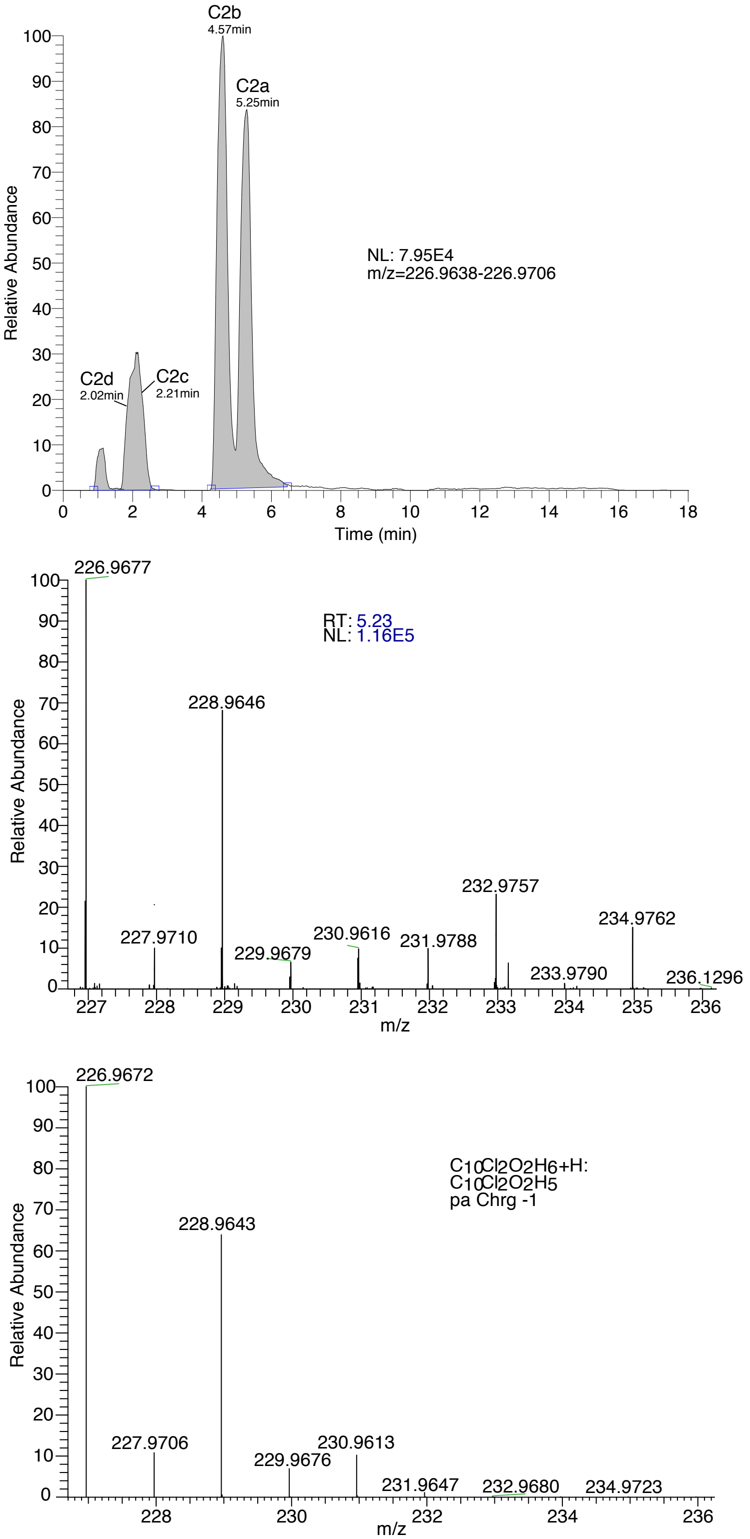


**Figure S6 I:** Extracted Ion Chromatogram, observed mass spectrum and theoretical mass spectrum for metabolites C2a-d m/z 226.9677 (carboxylated dichloroindene, loss of 8 Cl), data from sample GT20 June 29th 2018.

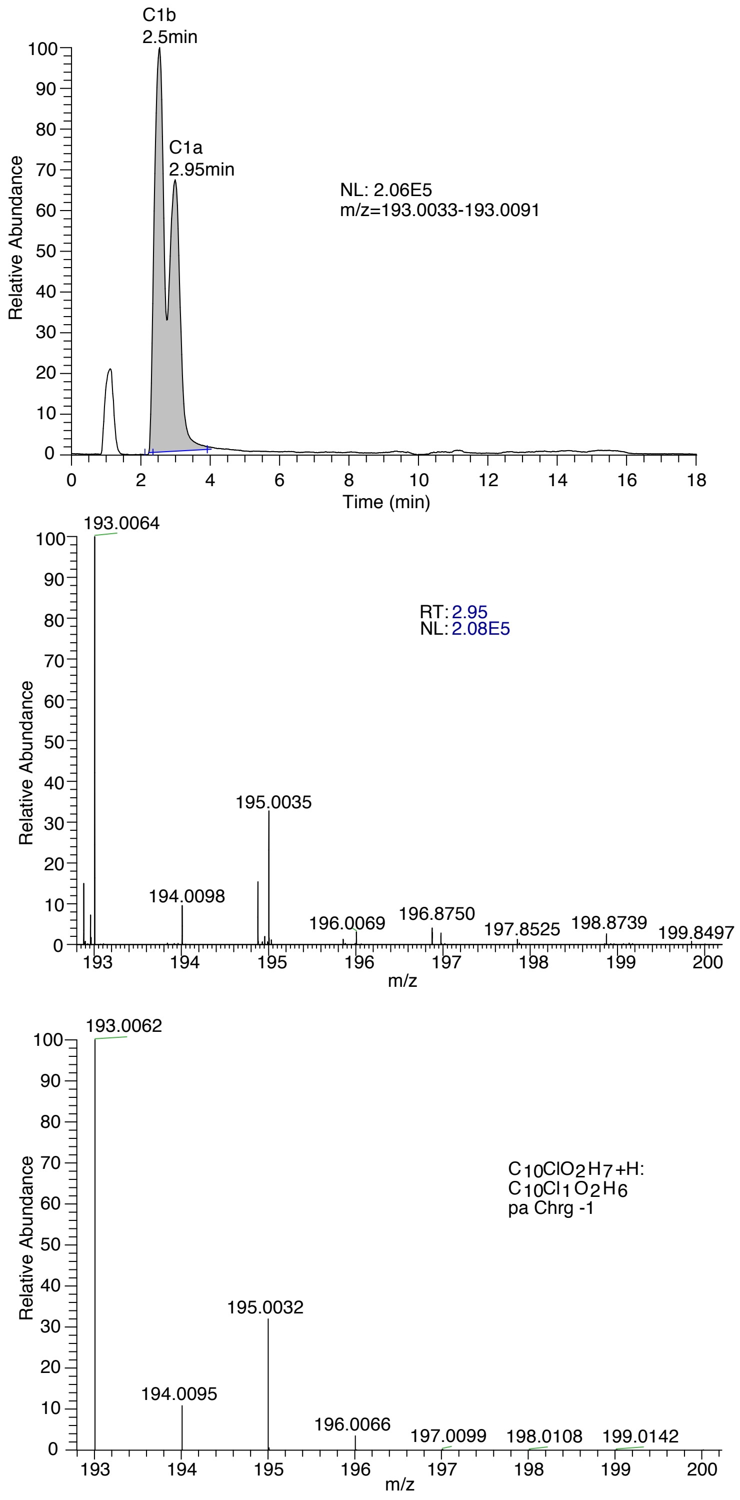


**Figure S6 J:** Extracted Ion Chromatogram, observed mass spectrum and theoretical mass spectrum for metabolites C1a-b m/z 193.0064 (carboxylated chloroindene, loss of 9 Cl), data from sample GT20 June 29th 2018.
